# Cold-water gut isolate from threespine stickleback *(Gasterosteus aculeatus)* reveals polypropylene surface oxidation and co-culture inhibition

**DOI:** 10.1101/2025.10.22.682630

**Authors:** Sarah M. Pasqualetti, Jolie Atwood, Abiodun Aderibigbe, Edward Russell, Kayleigh O’Keefe, Victoria Rosario, Kelly Ireland, Kenneth Sparks, Koral Campbell, Ruth Y. Isenberg, Ryan Lucas, Priyal Dhage, Kathryn Milligan-McClellan

## Abstract

Polyethylene terephthalate (PET) and polypropylene (PP), two of the most widely produced plastics in the United States, persist in cold-water environments where plastic-degrading microbes have been poorly characterized. Understanding how gut microbes interact and contribute to plastic degradation is essential for developing potential microbiome-based bioremediation strategies. We isolated 184 microbes from wild Alaskan threespine stickleback (*Gasterosteus aculeatus*) guts across six lakes and screened for plastic degrading potential using lipase/esterase assays and biofilm formation on PET and PP. During the screen for microbes with plastic degrading potential, we discovered that stickleback gut microbiota members enhance and suppressed the lipase, esterase, and biofilm activity of other microbes. Isolates with the highest plastic degrading potential were incubated in minimal media with PET or PP as the sole carbon source to determine whether plastic degradation potential is enhanced. Surface analysis identified a *Pseudomonas trivialis* strain that exhibited degradation of PP in monoculture; however, this activity was suppressed in the presence of another gut isolate, *Pseudomonas germanica*. These results demonstrate that microbes associated with the wild threespine stickleback gut microbiome possess plastic degradation potential and provide insights into how microbial interactions can either promote or inhibit bioremediation of plastic pollution in cold-water environments.

## Introduction

Plastic waste, such as polyethylene terephthalate (PET) and polypropylene (PP), is ubiquitous in aquatic ecosystems and degrades slowly^1–3^, fragmenting into micro- and nanoplastics that threaten biodiversity and ecosystem function. Low recycling rates^4^ and high landfill deposition exacerbate environmental loads, particularly in waters frequented by wildlife. The global scale of plastic pollution in aquatic environments threatens biodiversity and ecosystem health^5–8^, highlighting the need for more effective waste management, particularly in aquatic ecosystems where its impact is most significant^6,9–11^.

Microbial degradation has emerged as a promising approach towards mitigating plastic pollution, yet most characterized plastic-degrading microbes come from soil or temperate marine settings^12–17^, leaving cold freshwater systems underexplored. Several bacterial species have been shown to degrade plastic, such as *Ideonella sakaiensis* 201-F6, which utilizes PET as a carbon source and produces the enzymes required for degradation^18^, as well as multiple *Bacillus* and *Pseudomonas* species with demonstrated plastic-degrading abilities^15,19–22^.

Moreover, many natural degradation processes involve cooperative interactions among microbial species. Co-culturing microbes has been shown to enhance plastic degradation^15,22,23^, evidenced by enhanced enzyme activity and detectable surface modifications to plastic samples. These synergistic effects likely stem from complementary metabolic processes such as cross-feeding or genetic co-regulation associated with degradation pathways^24,25^. However, microbial interactions are not always synergistic. Antagonism between species can suppress degradation potential, which is a dynamic that is poorly studied in microorganisms in cold-, freshwater environments. Understanding both cooperative and inhibitory dynamics is therefore essential in identifying potential plastic degrading microorganisms in underexplored environments.

Given the global scale of plastic pollution, there is a need for accessible and relatively cheap screening methods to assess microbial plastic degradation potential. Similar large-scale screens have been conducted in other studies aiming to identify plastic-degrading organisms^26,27^.

However, these have rarely targeted isolates from organisms living in cold-, freshwater ecosystems, highlighting an important gap in knowledge.

To better understand plastic degradation, it is important to outline the essential microbial processes in biodegradation. Plastic degradation begins with the formation of biofilm on the plastic surface. Next, the microbial community secretes extracellular enzymes such as laccases^28^, lipases, esterases, or cutinases^29^. Lipases and esterases are commonly associated with polymer degradation due to lipid and polymer chain structure similarities. Hydrolytic enzymes^30–34^ cleave the long carbon backbone through hydrolysis, and the resulting subunits separate from the original molecule and are used by microbes in metabolic processes such as cellular respiration, where the final product of this degradation process is the release of carbon dioxide and water^18^.

To evaluate the ecological relevance of plastic degradation in cold-water environments, we explored host associated microbes in the threespine stickleback fish *(Gasterosteus aculeatus)*, a cold-water species found across the northern hemisphere in marine and freshwater sources^35^, most of which are plastic polluted^36^. The stickleback gut microbiome is diverse across populations, reflecting the wide range of environments they inhabit^37^. Diet plays a key role in shaping gut microbial diversity^38^, and the presence of environmental pollution could contribute to the presence of potential plastic degrading enzymes^17^. Marine and freshwater populations of stickleback share similar gut-associated microbes that include *Pseudomonas* and *Bacillus* species^39^, which could indicate an ability of the stickleback gut to degrade plastic. Therefore, using microbes from wild caught fish provide ecologically relevant results on how different wild populations respond to contaminant exposure.

Here, we asked whether microbes isolated from wild Alaskan threespine stickleback guts had the potential to degrade plastic. To answer this, we isolated 184 microbes from wild Alaskan threespine stickleback fish from six lakes and characterized plastic degrading potential by determining lipase and esterase activities, biofilm formation on PET and PP films, and screening for potential PP and PET degradation using attenuated total reflectance Fourier transform infrared spectroscopy (ATR-FTIR) and x-ray photoelectron spectroscopy (XPS). Surface modifications were observed using scanning electron microscopy (SEM). In addition to individual isolates, we also tested co-cultures to evaluate whether microbial interactions could enhance or inhibit plastic degradation. We hypothesized microbes with positive enzyme activity, biofilm formation, and detectable surface modifications on plastic films would have the greatest plastic degradation potential, and that co-cultures might improve these effects through synergistic interactions.

### Experimental Procedures

#### Microbe Isolation

The microbes tested in this study were sourced and isolated from healthy adult threespine stickleback guts from four freshwater and two anadromous populations in Southcentral Alaska (Table 1) under University of Alaska Anchorage IACUC Protocol 907288 and Alaska Department of Fish and Game (ADF&G) permit number P16-003, as well as University of Connecticut IACUC Protocol A21-021 and ADF&G permit number P-21-012.

**Table 1.** Characteristics of lakes from which microbes were isolated. Human activity was determined based on recreational activity, housing density, and location.

| Lake Name | Collection Coordinates | Elevation (m) | Area (ha) | Maximum Depth (m) | Human Activity |
| --- | --- | --- | --- | --- | --- |
| Bear Paw Lake<sup>41</sup> | 61.613948,<br>-149.752637 | 82 | 18.2 | 5.2 | Low |
| Big Lake<sup>41</sup> | 61.54787,<br>-149.85357 | 43 | 1009.7 | 27.1 | High |
| Cheney Lake<sup>41,42</sup> | 61.20490,<br>-149.75940 | 61 | 9.9 | 4.9 | High |
| Mud Lake<sup>41</sup> | 61.5625,<br>-148.9486 | 791 | 16.2 | 2.4 | Low |
| Rabbit Slough<sup>43</sup> | 61.53666,<br>-149.2533 | 13 | 8 km Shallow Creek | Unspecified | Moderate |
| Westchester Lagoon<sup>41,43,44</sup> | 61.20622,<br>-149.92423 | 4 | 33 | 4 | High |

Microbes from dissected guts were isolated as described in Milligan-McClellan et al. (2016)^40^. Microbes were streaked onto LB agar from original freezer stocks and incubated at 25°C for 24-48 hours to promote faster growth.

#### Microbe Identification

Microbes were identified by amplification and Sanger sequencing of the 16S or 18S ribosomal RNA gene using the 27F and 1492R universal primers^41^ for 16S or the Euk1A-F and 564R universal primers^42^ for 18S (Genewiz, New Jersey). Aligned consensus sequences were compared to NCBI’s core nucleotide database using the Basic Local Alignment Search Tool for nucleotides (BLASTn)^43^.

#### Culture Conditions

Microbes were streaked for isolation onto LB agar from original freezer stocks. Isolates were grown with shaking to mid-log phase at 25°C in LB and diluted to an OD_600_ of 0.5 for the lipase, esterase, and biofilm assays. Strains 9.1 (*Bacillus thuringiensis C15*) and 9.2 (P*seudomonas sp. B10),* both provided by Jay Mellies, are part of a consortia reported to degrade PET^15^ and were used as positive controls.

### Lipase Assay

#### Qualitative Plate Screening

Lipase activity of individual isolates was first screened on Rhodamine B agar plates. The agar was prepared following the Alken Murray Corporation^44^, modifying the procedure by emulsifying the lipids in a Conair™ Waring™ Laboratory Blender (22,000 rpm, 5 min). Plate inoculation was performed as described by Roberts et al. (2020)^15^ with the following modifications. A 5 µL aliquot of diluted culture was spotted onto triplicate Rhodamine B agar and incubated at 25°C. The formation of fluorescent orange colonies under UV illumination (302nm) indicates enzymatic hydrolysis of lipids, and the presence of this fluorescence was recorded at 24 and 48 hours to identify lipase-positive strains^45^.

#### Quantitative Co-culture Assay

For co-cultures, a quantitative liquid lipase assay was adapted from Zottig et al. (2016)^46^. A liquid media was adapted from the Alken Murray Corporation^44^, containing 0.9% (w/v) nutrient broth and 0.25% (w/v) yeast extract, and 0.02% (w/v) Rhodamine B. Assays were conducted in black, clear-bottom 96-well plates (Corning^®^, Cat: 3603) containing 190 µL of medium. Wells were inoculated with 10 µL of a single isolate (monoculture) or 5 µL each strain of two isolates (co-cultures). Culture plates were incubated without shaking at 25°C for 48 hours.

#### Fluorescence Measurement and Statistical Analysis

Fluorescence was measured at 0, 24, and 48 hours on a SpectraMax® i3x plate reader (350nm excitation and 580nm emission)^46^. Prior to each top-read measurement, plates were shaken for 5 seconds at medium speed to ensure homogeneity. All statistical analyses were performed in Prism 10. The effect of co-culturing was quantified by calculating the log fold change in fluorescence relative to the corresponding monocultures. One-sample t-tests were then used to determine if these log fold changes were significantly different from zero.

### Esterase Assay

1% and 2% tributyrin agar was prepared as described by Ramnath et al. (2017)^47^, but includes the following modifications: Agar plates were perforated using a 6mm biopsy punch. For monoculture assays, 20 µl of diluted culture was added to each punch-out. For co-culture assays, wells received either 20 µL of a single stain of a single isolate (monoculture) or 10 µL of each strain of two isolates (total 20 µL). Plates were incubated at 25°C for 48 hours. Esterase activity was inferred from clearing zones produced by tributyrin hydrolysis in the otherwise opaque medium. The diameter of each clearing zone (edge-to-edge, inclusive of the punch) was measured at 24 and 48 h. Equation (1) demonstrates the calculation for Total Hydrolysis Area (A_Hydrolysis_) below:

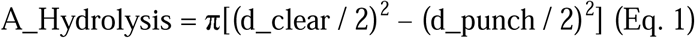

where d_clear_ is the clearing zone diameter and d_punch_ = 6 mm.

#### Statistical Analysis

All statistical analysis was performed in Prism 10. Monoculture samples were compared to positive control 9.1 *Bacillus thuringiensis* using a one-way ANOVA. For co-culture assays, log_2_ fold-changes in hydrolysis area were computed relative to the corresponding monoculture(s), and one-sample t-tests were used to assess whether log_2_ fold-changes differed significantly from zero.

### Biofilm Formation Assay

#### Preparation of Surfaces and Assay Setup

Polypropylene (PP; Millipore Sigma, Cat. GF99473552) and polyethylene terephthalate (PET; Millipore Sigma, Cat. GF31588229) films were cut into 1×1 cm squares and sterilized in 70% ethanol. Before use, each film was aseptically handled with sterile forceps, rinsed first in 96% ethanol, then in sterile double-distilled water (ddH_2_O). The assay was performed in 48-well polystyrene culture plates (Fisher Scientific, Cat. 08-772-52) containing LB. The experimental treatments, each performed in triplicate, consisted of wells containing: (1) a vertically oriented PET film, (2) a vertically oriented PP film, or (3) no film (control). Microbes were either added to the culture plate directly from an agar plate (initial screen) or grown in an overnight culture with shaking at 25°C (quantified and reported assay) at a final OD_600_ of 0.002. Plates were incubated at 25°C without shaking for 68 hours.

#### Biofilm Staining and Quantification

Following incubation, biofilms formed on the plastic films and on the surfaces of the polystyrene wells were quantified separately. Plastic films were aseptically transferred to a new 48-well plate for analysis. Both the original plate (containing well-adhered biofilms) and the new plate (containing film-adhered biofilms) were processed for staining. Biofilm fixation and staining were performed as described in Chen et al., 2022^48^.

Crystal violet was aspirated from the wells and films and rinsed in deionized water (diH_2_O). All samples were dried at 30°C overnight. Crystal violet was extracted with 30% (v/v) glacial acetic acid (Sigma, Cat. A6283) and the absorbance was measured at 590nm.

#### Data Normalization

All statistical analyses were conducted in Prism 10. The relative biofilm formation (RBF) for each replicate was calculated as the ratio of its absorbance value to that of the blank control; see Eq. (2) below:

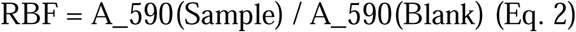

Relative biofilm formation for each condition was then normalized (RBF_Normalized_) to the total surface area and the respective control to account for residual crystal violet; see Eq. (3) below:

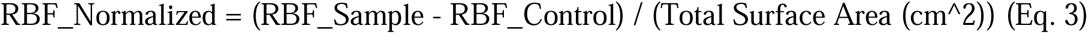

To compare biofilm growth on different surfaces, a two-way ANOVA with Dunnett’s multiple comparisons test was used to analyze differences between the PET, PP, and no-plastic control conditions^49^. To assess co-culture interactions, a one-way ANOVA with Dunnett’s test was used to compare biofilm formation in co-cultures to their respective monocultures ^49^. Additionally, the log fold change in relative biofilm formation for co-cultures was calculated, and one-sample t-tests were used to determine if these changes were significantly different from zero.

### Growth for Biofilm Assays, Fourier Transform Infrared Spectroscopy, and X-Ray Photoelectron Spectroscopy

Minimal salt media (MSM) was modified from Augusta et al., 1993^50^ and contained 0.1% (w/v) NH_4_NO_3_, 0.07% (w/v) KH_2_PO_4_, 0.07% (w/v) K_2_HPO_4_, 0.07% (w/v) MgSO_4*_7H_2_0, 0.5% (v/v) Wolfe’s Trace Vitamins, and 0.5% (v/v) Trace Minerals solutions. Liquid cultures were grown to mid-log phase in LB, pelleted at 8000 x g for 10 minutes, and resuspended in MSM to an OD_600_ of 0.5. Diluted cultures were added to a glass tube containing 4 mL MSM and 1×1cm PP or PET films as a carbon source. Monocultures were inoculated with 1 mL of a single strain while co-cultures were inoculated with 500µL of each strain and incubated at 25°C for 8 weeks under static conditions. To maintain nutrient availability, fresh media was added to each tube at the 4-week midpoint.

At the end of the incubation period, the plastic films were removed and divided into half for use in biofilm, FTIR, and XPS.

### Attenuated Total Reflectance Fourier Transform Infrared Spectroscopy (ATR-FTIR)

Plastic films were treated with enzymatic lysis buffer containing 20 mg/mL lysozyme overnight at 37°C to remove the cells and biofilm, followed by three rinses with ddH_2_O water and drying at 30°C.

ATR-FTIR spectra were collected according to Kyaw et al. (2012) and Roberts et al. (2020)^15,21^ with modifications. Spectra were collected on a Nicolet™ iS20 FTIR spectrometer using OMNIC^TM^ software (Thermo Fisher Scientific) over a range of 4000 to 400 cm^-1^ at a resolution of 8 cm^-1^ and a total of 16 scans per collection. Each plastic film was measured in triplicate (*n*=9 total), and the averages were compared both to uninoculated controls and an untreated blank. The carbonyl index of PP was calculated using the specified area under band (SAUB) method^51^, shown in Eq. 4:

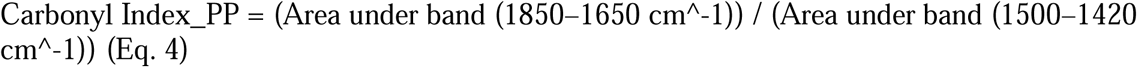

The carbonyl index of the PET was calculated according to Roberts et al., 2020^15^, shown in Eq. 5:

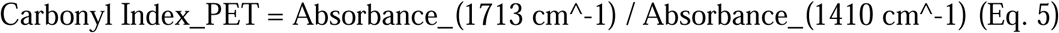

Statistical analysis was performed in Prism 10 and compared to a negative control using a one-way ANOVA.

### X-Ray Photoelectron Spectroscopy (XPS)

XPS was performed according to Yang et al. (2014)^52^ with modifications. XPS was recorded on a ThermoFisher Scientific K-Alpha spectrometer (Part no.1200505) equipped with a monochromatic Al Kα source (hv=1486.6eV). Samples were mounted to the sample holder using double sided carbon tape. The X-ray spot size was 400 µm, take off angle was 90°, and the analyzer chamber pressure was maintained below 5.0×10^-8^ mbar before scans were recorded.

Triplicate scans were performed on each plastic film. Surface charging was mitigated using an electron flood gun. Survey scans were collected with a pass energy of 200 eV and an energy step of 1.0 eV, while core scans were acquired at 50 eV with a 0.1 eV step size. The dwell time was 50 ms per point. All spectra and peak fittings were carried out according to Beamson and Briggs (1992)^53^ using CasaXPS and charge correction was done with reference to C 1s photoelectron line at 285 eV.

Statistical analysis was performed in Prism 10 to compare peak area percentages among chemical groups using a two-way ANOVA followed by Tukey’s multiple comparisons test^54^. To evaluate differences in overall XPS peak distributions across treatment groups, pairwise permutational multivariate analysis of variance (PERMANOVA) was performed in R^55^ (v4.5.1) using the Vegan package^56^ (v2.7-1). Bray-Curtis dissimilarities were calculated from peak area percentages, and significance was assessed with 9999 permutations.

### Scanning Electron Microscopy (SEM)

The same plastic films analyzed by FTIR were examined for degradation using SEM. Sample preparation and imaging were adapted from Jeon et al. (2021)^20^. Samples were mounted vertically onto double-sided carbon tape on a variable-tilt specimen holder and sputter-coated with a 5 nm layer of Au-Pd (80:20) using a Safematic CCU-010 sputter coater. Imaging was performed at 90° with an FEI Nova NanoSEM 450 equipped with an Everhart-Thornley Detector (ETD) under high vacuum conditions. SEM was performed at the University of Connecticut’s Bioscience Electron Microscopy Laboratory CORE facility.

## Results

### Identification of stickleback gut isolates

Identification for all organisms is available in Supplementary Data Table 1. Given the large dataset, a subset of isolates will be highlighted in the results for lipase, esterase, and biofilm forming abilities in monoculture and co-culture (Table 2). The full results of all 184 isolates are in Supplemental Table 1.

**Table 2.** Identifications of the representative subset of microbes highlighted in the results. Lipase and esterase activity results were based on plate-based assays.

| Isolate | Species | Lipase Activity | Esterase Activity |
| --- | --- | --- | --- |
| KMM 191 | *Pseudomonas germanica* | + | + |
| KMM 193 | *Pseudomonas fragi* | + | + |
| KMM 195 | *Pseudomonas trivialis* | + | + |
| KMM 200 | *Erwinia persicina* | - | - |
| KMM 202 | *Pantoea agglomerans* | - | + |
| KMM 230 | *Pseudomonas germanica* | + | + |

### Stickleback gut isolates have variable and co-culture specific lipase activity

In total, 74 of the 184 isolates were positive for lipase activity (Supp. Table 1). There were no populations for which all isolates were lipase positive, nor genera of microbes in which all isolates were lipase positive. For example, of the 15 *Pseudomonas* isolates tested, 9 were positive. There were also differences within species such as *Aeromonas salmonicida* strains KMM 197 (positive) and KMM 284 (negative) (Supp. Table 1). Among the positive isolates, KMM 191 (2.1×10^6^ ± 4.0×10^5^ RFU) and KMM 230 (1.8×10^6^ ± 7.9×10^5^ RFU), both *Pseudomonas germanica,* demonstrated the highest lipase activity in monoculture whereas other strains such as KMM 202 (*Pantoea agglomerans)*, showed lower lipase activity (5.4×10^5^ ± 2.4×10^5^ RFU) (Figure 1A).

**Figure 1.**
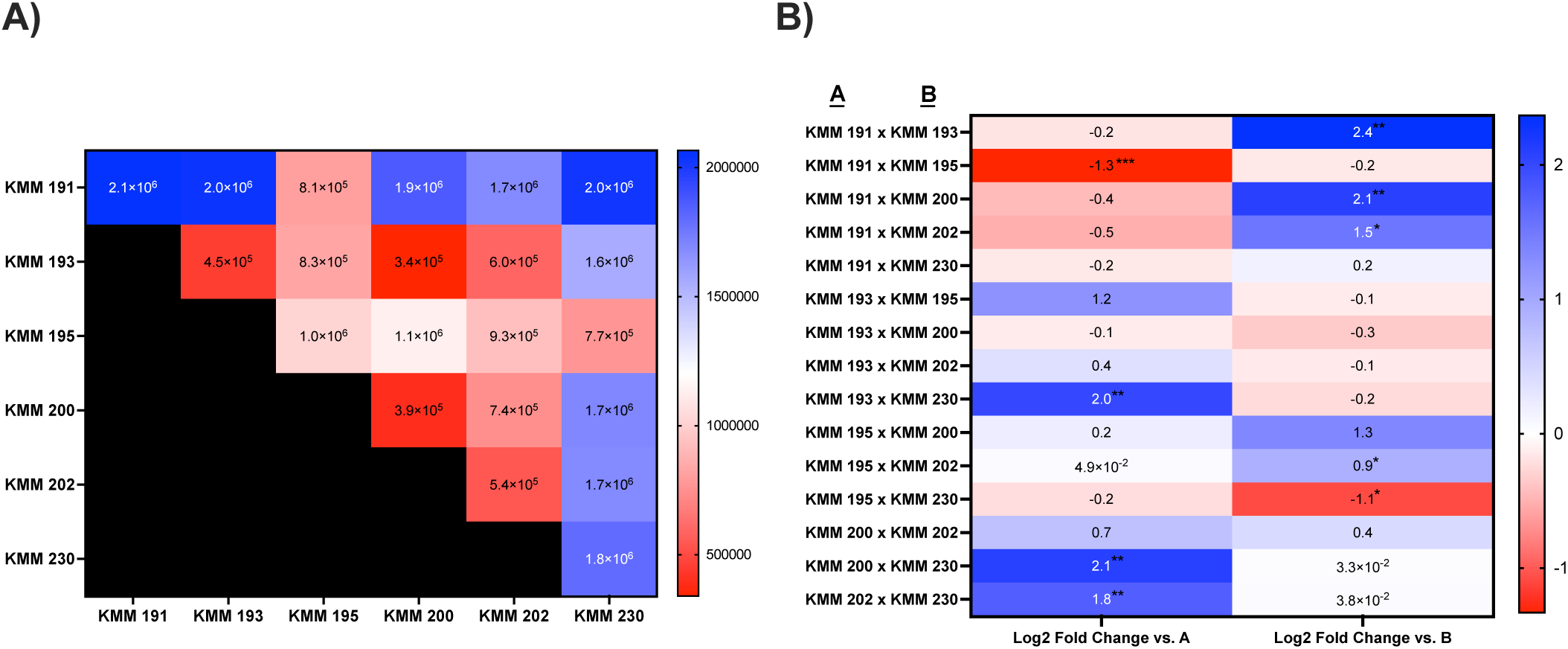
Lipase activity and fold change analysis in monoculture and co-culture. **A)** Heat map of lipase activity (reported as RFU) for each monoculture and co-culture. Black cells are repeated comparisons. **B)** Log_2_ fold change in lipase activity for each co-culture relative to monocultures of microbe A (right) and microbe B (left). Positive values indicate increased lipase activity in co-culture, and negative values indicate decreased activity. Statistical significance is indicated by p < 0.05 (*) and p < 0.01 (**).

Co-cultures had both enhanced and suppressed lipase activity relative to the monocultures (Figure 1B). Co-culture of KMM 195 (*Pseudomonas trivialis)* and KMM 200 (*Erwinia persicina*) had increased lipase activity compared to individual isolates with log_2_ fold increases of 0.2 vs. KMM 195 and 1.3 vs. KMM 200. A similar trend was observed in the co-culture of KMM 200 and KMM 202 had increased lipase activity compared to individual isolates with log_2_ fold increases of 0.7 vs. KMM 200 and 0.4 vs. KMM 202. In contrast, co-culture of KMM 191 and KMM 195 resulted in a -1.3 log_2_ fold decrease compared to monoculture of KMM 191 and a -0.2 log_2_ fold decrease compared to KMM 195, suggesting suppression of lipase activity of both strains. Similarly, co-culture of KMM 195 and KMM 230 (*P. germanica)* resulted in a -0.2 log_2_ fold decrease vs. KMM 195 and -1.1 log_2_ decrease vs. KMM 230. These results highlight that lipase activity in co-culture is highly variable and dependent on specific strain pairings.

### Species, population origin, and co-culture interactions influence esterase activity among stickleback gut isolates

Hydrolysis area on tributyrin agar directly correlates with the amount of esterase activity^47,57^, thus higher esterase activity would suggest higher plastic degrading potential^58^. Isolates were tested on both 1% and 2% tributyrin agar, as esterase activity can vary with substrate concentration^47^. While some isolates were positive on only one concentration, the differences were negligible, and overall results were consistent across both conditions for all isolates. While 138 of the 184 isolates were positive for esterase activity (Supp. Table 1), 21 of those microbes had at least 1.6 times higher esterase activity (*p* < 0.05) than positive control 9.1 (104.4 ± 0.0). The total hydrolysis area of the isolates with the largest clearing zones exceeded 173.3mm^2^ (Supp. Table 2).

Out of these 21 isolates, all were *Aeromonas* or *Pseudomonas* species apart from *Chromobacterium aquaticum (*KMM 293, 173.3 ± 25.2 mm^2^). KMM 191 (*P. germanica)* had the highest esterase activity of all the isolates, followed by KMM 195 (*P. trivialis*).

Notably, 17 of the 21 microbes were isolated from fish living in lakes with moderate to high human activity, suggesting human activity could contribute to the plastic degrading potential of a microbiome. Among the isolates with the highest esterase activity, 11 of the 21 isolates were *A. veronii* from Rabbit Slough, with hydrolysis areas ranging from 181.4 to 276.7 mm^2^ (Supp. Table 2). Similar to the lipase activity, there were other *A. veronii* strains, some isolated from the same population of stickleback, with lower esterase activity, indicating within species and within population variation in esterase activity. For example, two *A. veronii* strains, KMM 358 and KMM 364, both from Rabbit Slough, have significant differences in esterase activity (118.6 ± 12.2 mm^2^ and 217.0 ± 15.9 mm^2^, respectively). Other species, such as *P. germanica* strains KMM 191 and KMM 230, had similar esterase activity (299.8 ± 81.4 mm^2^ and 227.77 ± 47.6 mm^2^, respectively) (Figure 2).

**Figure 2.**
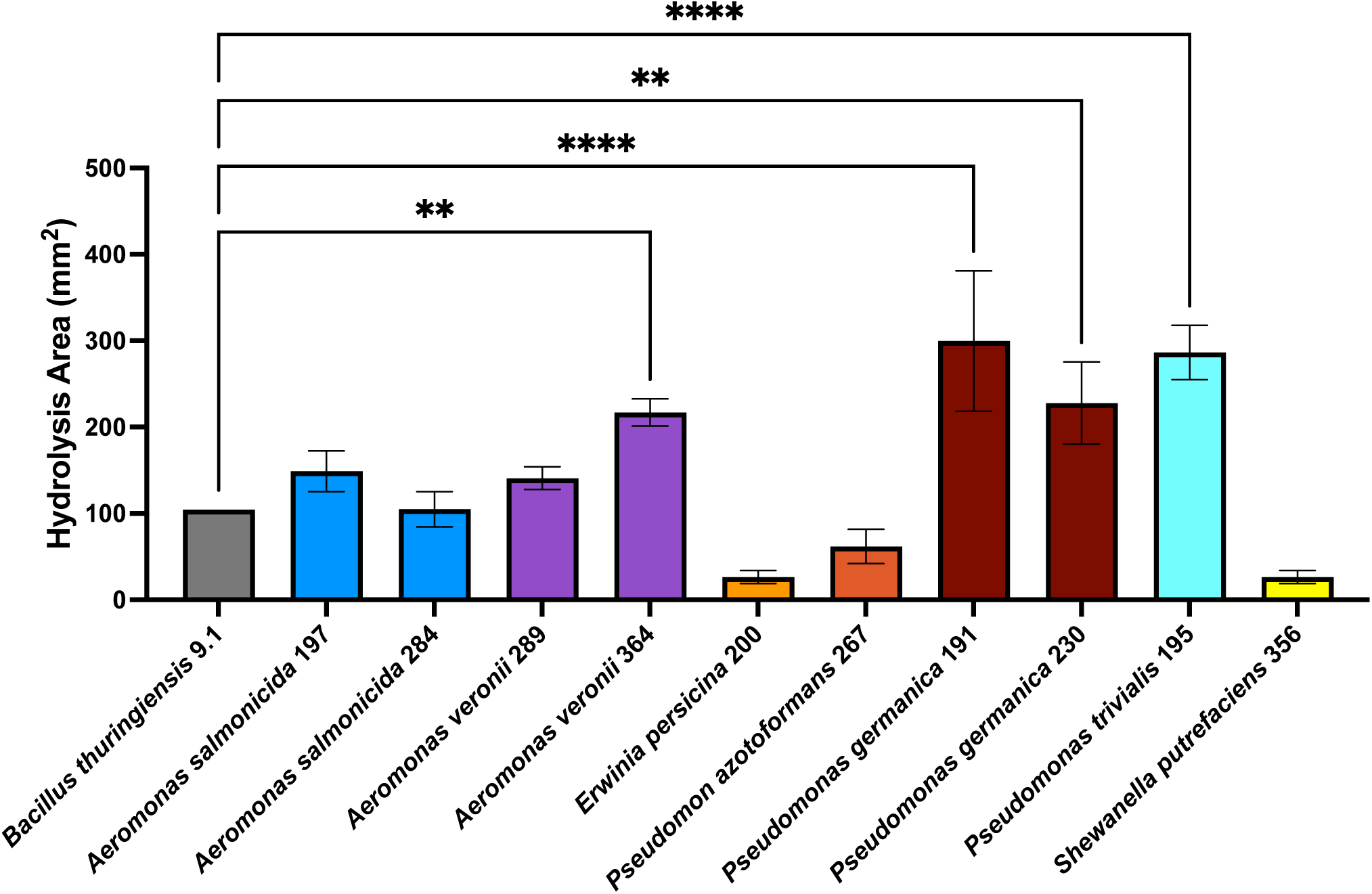
Esterase activity of stickleback-derived isolates after 48 hours on 1% tributyrin agar. Esterase activity was quantified and compared to positive control 9.1^15^. Statistical significance is indicated by p < 0.002 (**), p < 0.0001(****). Error bars indicate standard deviation.

All isolates with significant esterase activity were also positive for lipase activity.

However, lipase activity did not predict esterase activity. For example, KMM 197 (*A. salmonicida,* 149.0 ± 23.55 mm^2^) and KMM 267 (*Pseudomonas azotoformans,* 61.78 ± 20.2 mm^2^) produced lipase without significant esterase activity. KMM 200 (*E. persicina,* 26.4 ± 7.7 mm^2^) and KMM 284 (*A. salmonicida,* 105.0 ± 20.5 mm^2^) did not have significant esterase activity and were not positive for lipase activity.

Co-cultures were evaluated for esterase activity and compared to corresponding monocultures. There were some co-cultures that demonstrated significant changes in esterase activity relative to the monoculture of an individual strain, however there were no co-cultures that showed significant increases or decreases relative to both monocultures (Figure 3A). For example, the co-culture between KMM 191 and KMM 193 (*Pseudomonas fragi*) had a significant log_2_ increase in esterase activity vs. KMM 193 (2.7, *p* = 0.003), but was insignificant compared to KMM 191 (-0.1, *p =* 0.239) (Figure 3B). This suggests that while some co-cultures exhibit enhanced esterase activity relative to a single partner, the overall activity of the co-cultures does not significantly enhance or inhibit the activity of both contributing monocultures or that the esterase activity is due to one strain in the co-culture.

**Figure 3.**
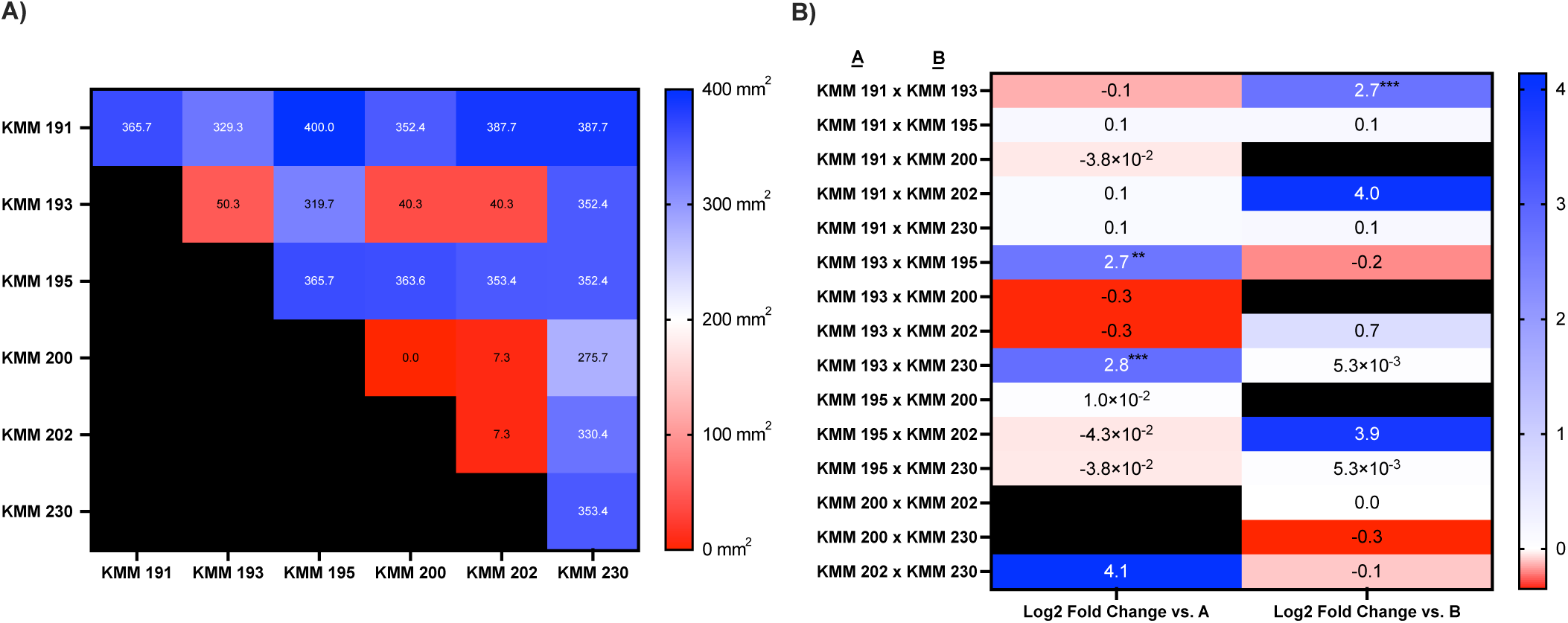
Esterase activity and fold change analysis in monoculture and co-culture using the same microbial isolates shown in Figure 1. **A)** Heat map showing esterase activity (reported as mm^2^) for each monoculture and co-culture. **B)** Log_2_ fold change in esterase activity for each co-culture relative to monoculture partners A (left) and B (right). Positive values indicate increased esterase activity, and negative values indicate decreased activity. Statistical significance is indicated by p < 0.01 (**) and p < 0.001 (**). Black cells represent repeated or non-applicable comparisons.

### Biofilm formation varies by plastic type and is modulated by co-culturing

While it is known that microbial biofilm formation on plastics contributes to adsorption, desorption, and degradation of plastics^59^, comparative studies on biofilm formation across different plastic types are limited. To address this, we asked whether the ability of the microbes to form biofilms depends on the type of plastic the microbes were exposed to. We initially screened all 184 microbes for the ability to form biofilms on plastics with cultures started directly from agar plates and then repeated the quantification of 61 microbes that had high biofilm formation in the initial screen with cultures started with a liquid culture to mid-log phase and standardized by surface area (cm^2^).

For all but two isolates, there was greater biofilm formation on the PS wells which accounted for the total surface associated biomass including the liquid-air interface, regardless of whether PET or PP films were added, compared to the biofilm formed directly on the PET or PP films. KMM 267 (*P. azotoformans*, 0.21 ± 0.1) and KMM 356 (*S. putrefaciens,* 0.17 ± 0.06) formed more biofilm on the plastic film compared to the PS well, in the presence of PET (0.1 ± 0.03, 0.07 ± 0.01 respectively) or PP (0.11 ± 0.06, 0.09 ± 0.01 respectively) (Figure 4). When there was no plastic film added, both isolates had the same relative biofilm formation of 0.06 ± 0.0 on the PS well (Figure 4).

**Figure 4.**
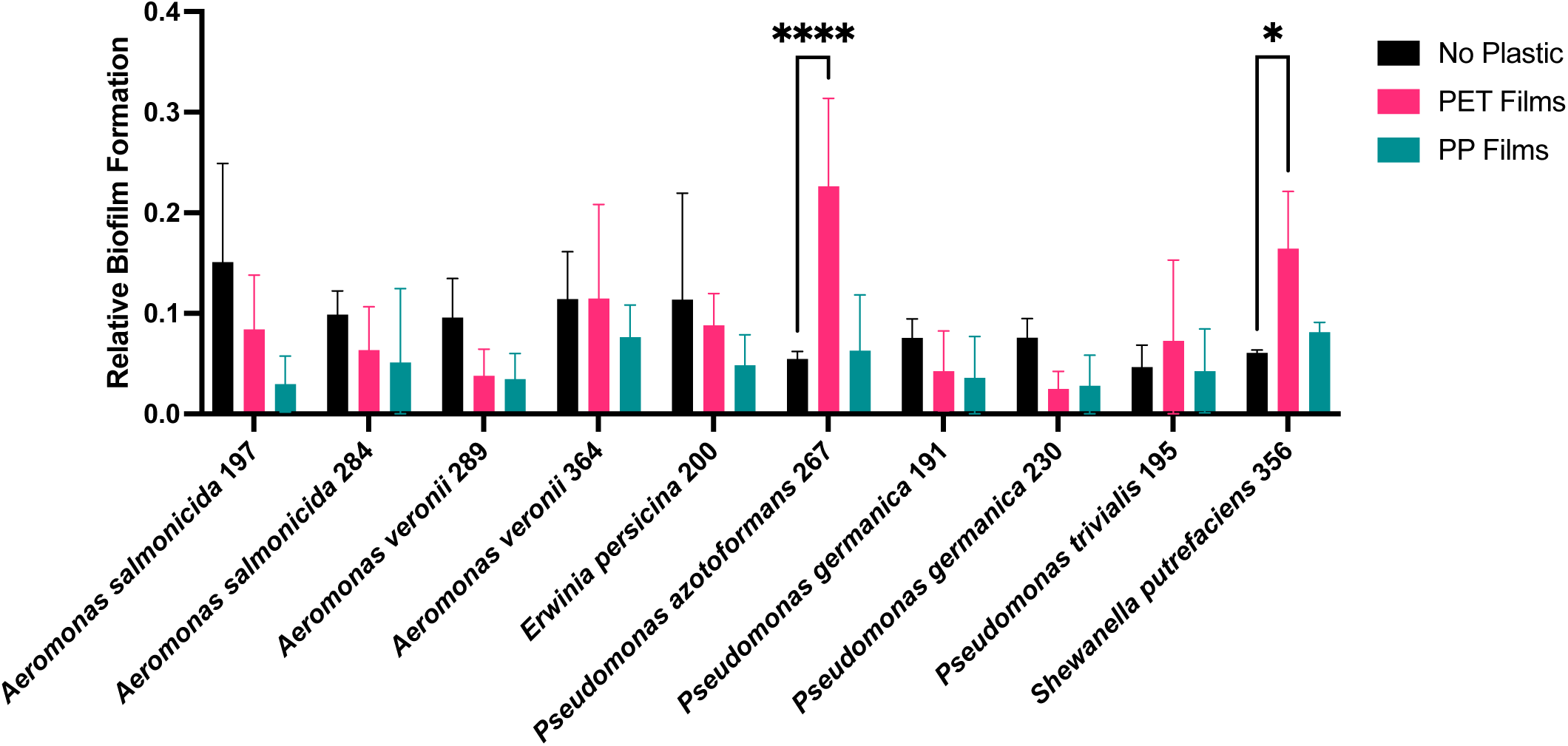
Comparison of biofilm formation on PET and PP films relative to control well containing no plastic film. Statistical significance is represented as p < 0.05 (*) and p < 0.001 (****). Error bars indicate standard deviation. Only statistically significant increases in biofilm formation on the plastic films or between plastic films are shown.

One isolate, KMM 267 (*P. azotoformans*), had significant differences in relative biofilm formation on the PET film (0.21 ± 0.09) compared to the PP film (0.06 ± 0.05), indicating a higher affinity for forming biofilms on the PET surface (Figure 4). In contrast, biofilm formation on the PP film was not significant (Figure 4).

To determine whether the presence of PET or PP contributed to increased biofilm formation, we quantified the total biofilm formed on the plastic film and the PS well walls and compared it to a control well without plastic film, hereby referred to as control (Figure 5A).

**Figure 5.**
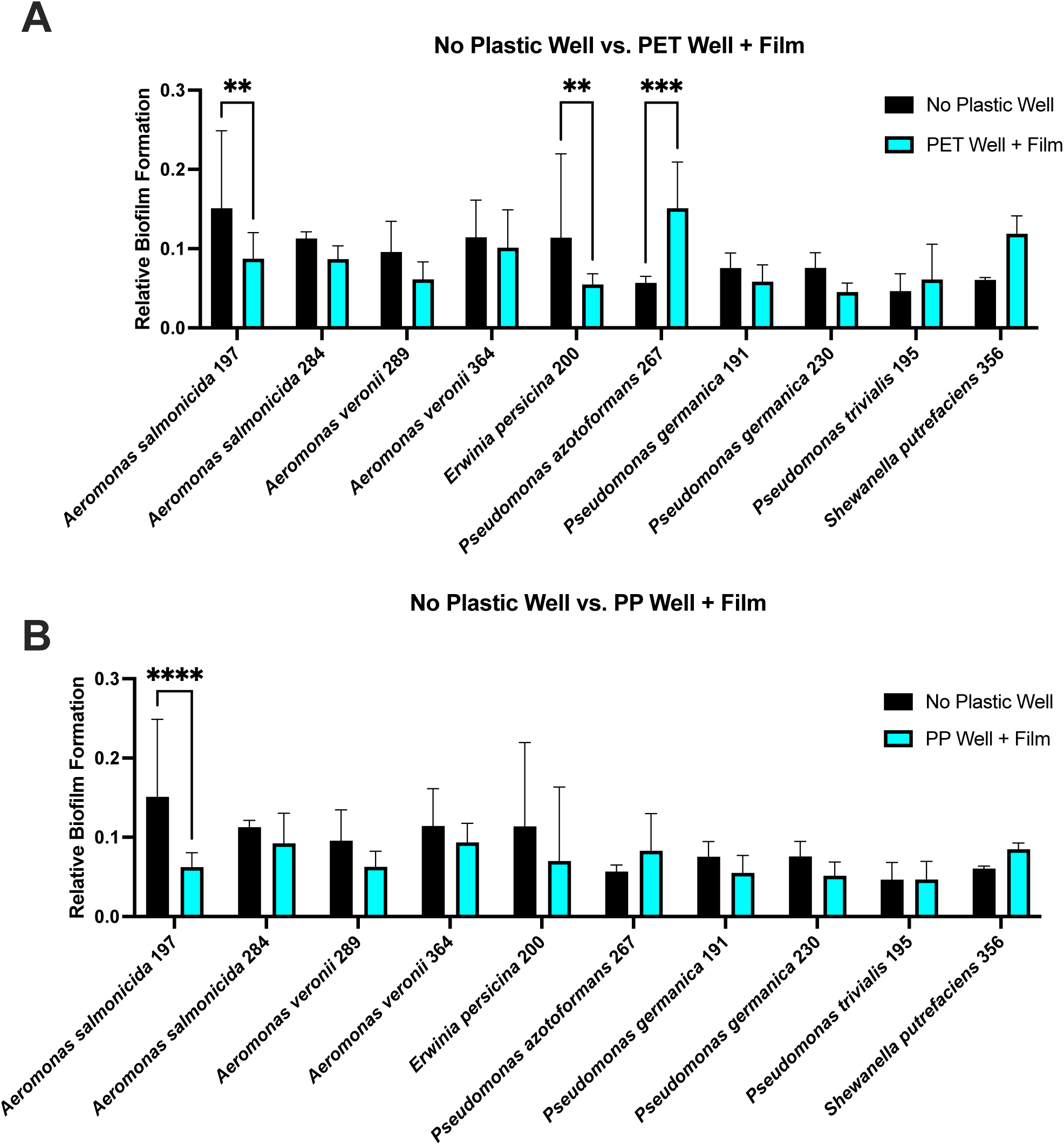
Total biofilm formation for PET (A) and PP films (B) and the incubation well was compared to wells containing no plastic film, showing differences in biofilm formation for several isolates. Statistical differences are represented as p values <0.003 (**), 0.001 (***), and <0.0001 (****). Error bars indicate standard deviation.

KMM 267 (*P. azotoformans*), formed more biofilm in the presence of PET (0.15 ± 0.06) than in the control (0.06 ± 0.01). Conversely, KMM 200 (*E. persicina*) showed reduced biofilm in the presence of PET or PP compared to control conditions without the addition of plastic films (0.114 ± 0.1) (Figure 5A-B). KMM 222 (*Pseudomonas peli*), formed more biofilm in the presence of PP (0.15 ± 0.1) compared to the control (0.05 ± 0.02), however there was no significant difference observed between the control and the well with PET added (Supp. Table 1).

KMM 197 (*A. salmonicida*) formed more biofilm in the control well (0.15 ± 0.1) compared to both treatment conditions with PET and PP added (Figure 5A-B). Other isolates not shown that had similar results included KMM 244 (*Buttiauxella ferragutiae*) and KMM 255 (*Lysinibacillus sphaericus*) (Supp. Table 1).

Relative biofilm formation was also measured for monocultures and co-cultures on both PP and PET films. Notably, the co-culture between KMM 191 and KMM 195 showed significantly inhibited biofilm formation on PP when in co-culture compared to monoculture with a -3.3 log_2_ fold decrease vs. KMM 191 and a -1.6 log_2_ fold decrease vs. KMM 195 (Figure 6A). Interestingly, the co-culture of KMM 191 and KMM 195 showed different results on PET, where there was a -0.7 log_2_ fold decrease vs. KMM 191 and a 0.1 log_2_ fold increase vs. KMM 195 (Figure 6B), which could suggest that biofilm formation is influenced by not only microbial interactions, but also the type of plastic surface.

**Figure 6.**
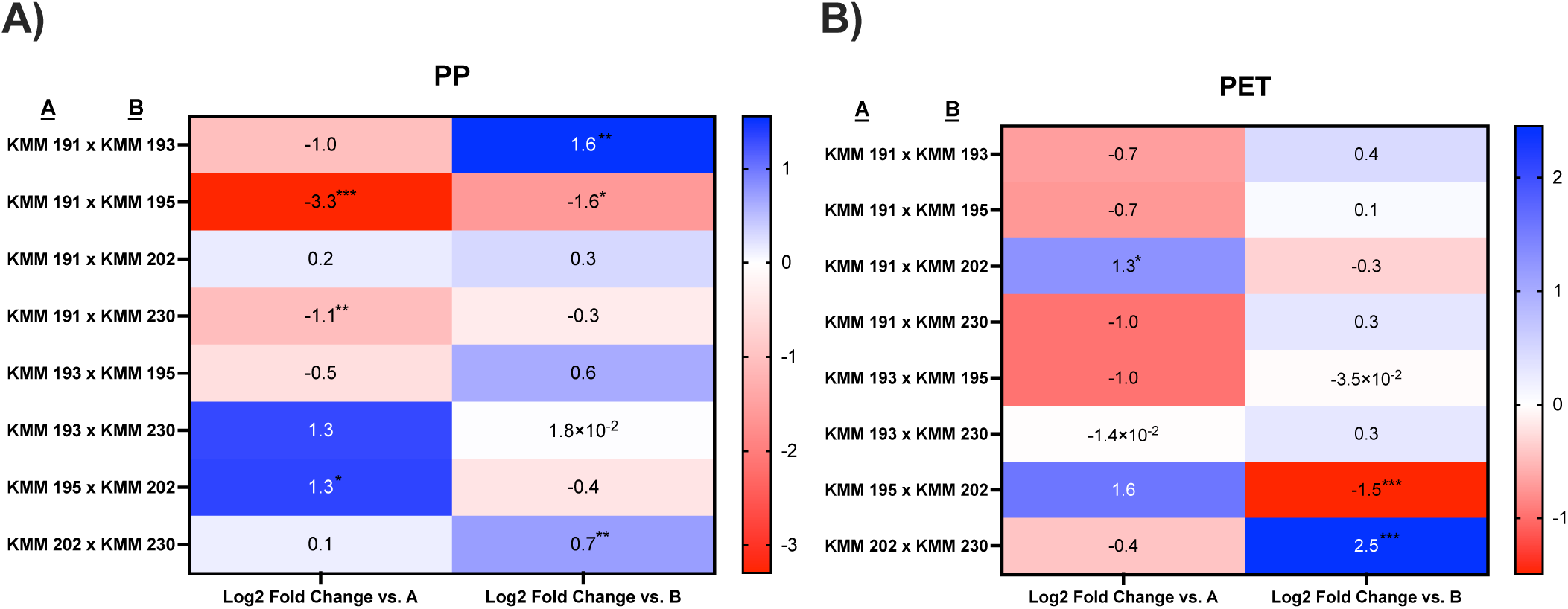
Log_2_ fold change in relative biofilm formation on PP (A) and PET (B) for each co-culture relative to monoculture partners A (left) and B (right). Positive values (blue) indicate increased biofilm formation and negative values (red) indicate decreased activity. Statistical significance is indicated by p values <0.05 (*), <0.003 (**), 0.001 (***).

### Surface characterization of polypropylene reveals isolate-specific oxidation that is inhibited in co-culture

To evaluate plastic degradation, ATR-FTIR was performed on 54 microbes on both PET and PP films. Microbes were chosen for FTIR analysis based on species, the presence or absence of lipase and esterase activity, and biofilm forming ability. This selection aimed to determine whether both enzyme production and biofilm formation were required for potential degradation. Out of the 54 microbes tested, no microbes were shown to individually degrade PET. Potential degradation was only observed on the PP films inoculated with KMM 195 (*P. trivialis)*, indicated by the formation of a carbonyl stretch in the spectra (Figure 7A). Carbonyl index calculations^51^ confirmed a significant increase in KMM 195 compared to the negative control (Figure 7B) across biological replicates, supporting the potential for plastic degradation. Notably, KMM 195 was the only *P. trivialis* out of the tested isolates and was isolated from Westchester Lagoon, a population with high human activity which may have contributed to its increased plastic degrading potential^17^.

**Figure 7.**
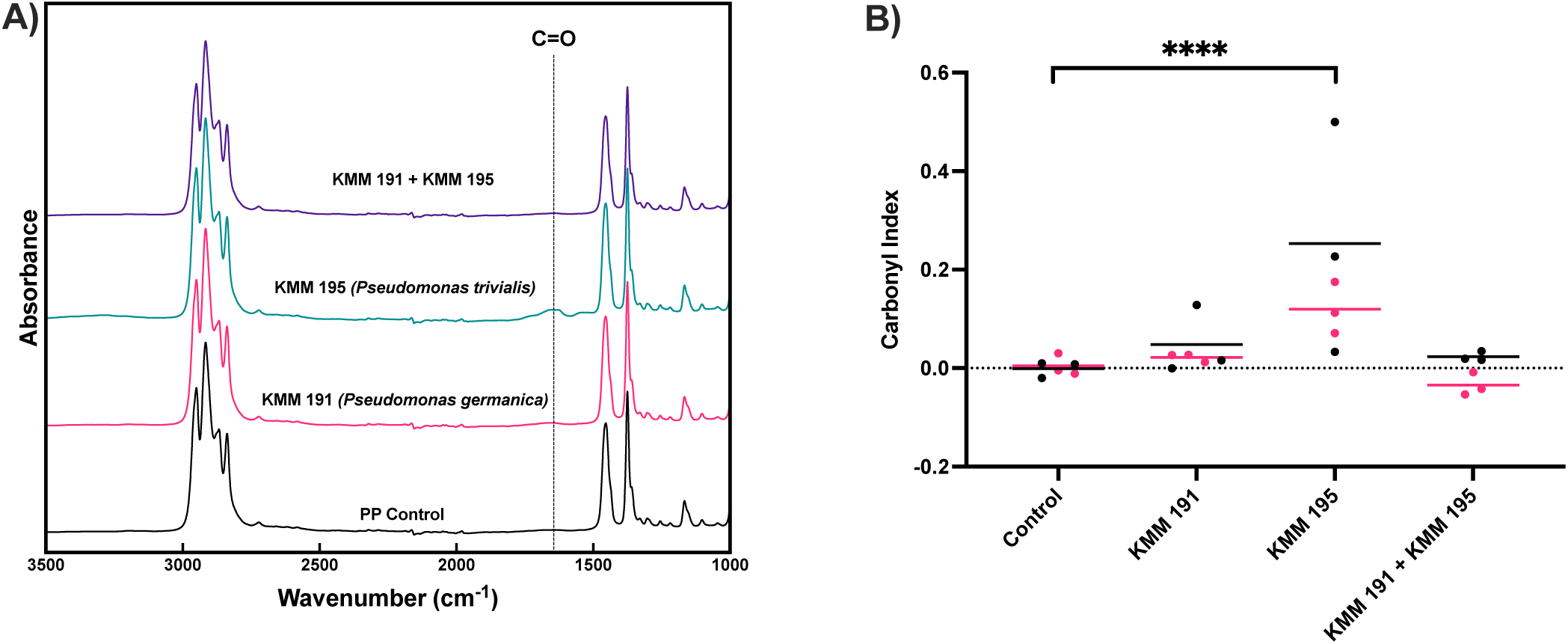
ATR-FTIR analysis revealed increased surface oxidation of PP films incubated with KMM 195, whereas co-culture with KMM 191 suppressed oxidative changes. **A)** Averaged ATR-FTIR spectra across biological replicates (*n*=27) of PP films from the control (black), KMM 191 (pink), KMM 195 (teal), and the KMM 191 + KMM 195 co-culture (purple). A dotted line highlights the emerging carbonyl peak in the KMM 195 monoculture. Inoculated samples were also compared to an untreated blank (not shown), which produced spectrum identical to the control. **B)** Comparison of carbonyl index of PP films inoculated with KMM 191 and KMM 195 as monocultures or as a co-culture. Each point represents a technical replicate (triplicate ATR-FTIR scans), with colors distinguishing biological replicates. Lines represent the mean for each biological replicate. Statistical significance was first assessed within each biological replicate, with only the KMM 195 treatment group significantly different from the control. The statistical significance shown (****p < 0.0001) reflects the combined analysis across both biological replicates.

Given that potential degradation was limited in monocultures, plastic films were incubated in co-cultures to determine whether interactions between strains could affect plastic degradation. Co-cultures for ATR-FTIR analysis were selected based on co-cultures with KMM 195 (*P. trivialis*) to determine whether potential degradation would be enhanced or suppressed. Notably, co-cultures of KMM 195 with KMM 191 (*P. germanica*) (Figure 7B), KMM 193 (*P. fragi)*, or KMM 230 (*P. germanica*) (data not shown) consistently resulted in a reduction in the carbonyl index, suggesting KMM 195 likely requires monoculture conditions for optimal plastic degradation potential on PP films.

Since these findings suggested that the co-culture between KMM 191 and KMM 195 suppressed degradation activity, X-ray photoelectron spectroscopy (XPS) was then performed on the PP films to identify the chemical composition of the polymer surface. XPS analysis showed significant shifts in surface chemical composition following incubation with KMM 195.

Representative C 1s spectra for the PP film incubated with KMM 195 (Figure 8A-B) showed a decrease in the C-C/C-H peak at 284.9eV and an increase in oxidized carbon species relative to the control. Peak area quantification (Figure 8D-G) confirmed that C-C/C-H was significantly reduced (p = 0.008) and C-O (286.7eV) was significantly elevated (p = 0.01). C=O (287.9eV) peak area increased with the KMM 195 treatment group and approached significance (p = 0.053). Comparisons of C-C/C-H, C-O, and C=O peak areas between KMM 195 and the KMM 191+ KMM 195 co-culture also trended toward significance (p = 0.061, 0.061, and 0.059, respectively). The increase in the relative abundance is considered suggestive of PP oxidation, further supporting the possibility KMM 195 has a potential to degrade PP in a monoculture. In contrast, PP films incubated with KMM 191 and KMM 195 showed no significant differences from the control (Figure 8C), further indicating that KMM 191 suppressed the degradative activity of KMM 195.

**Figure 8.**
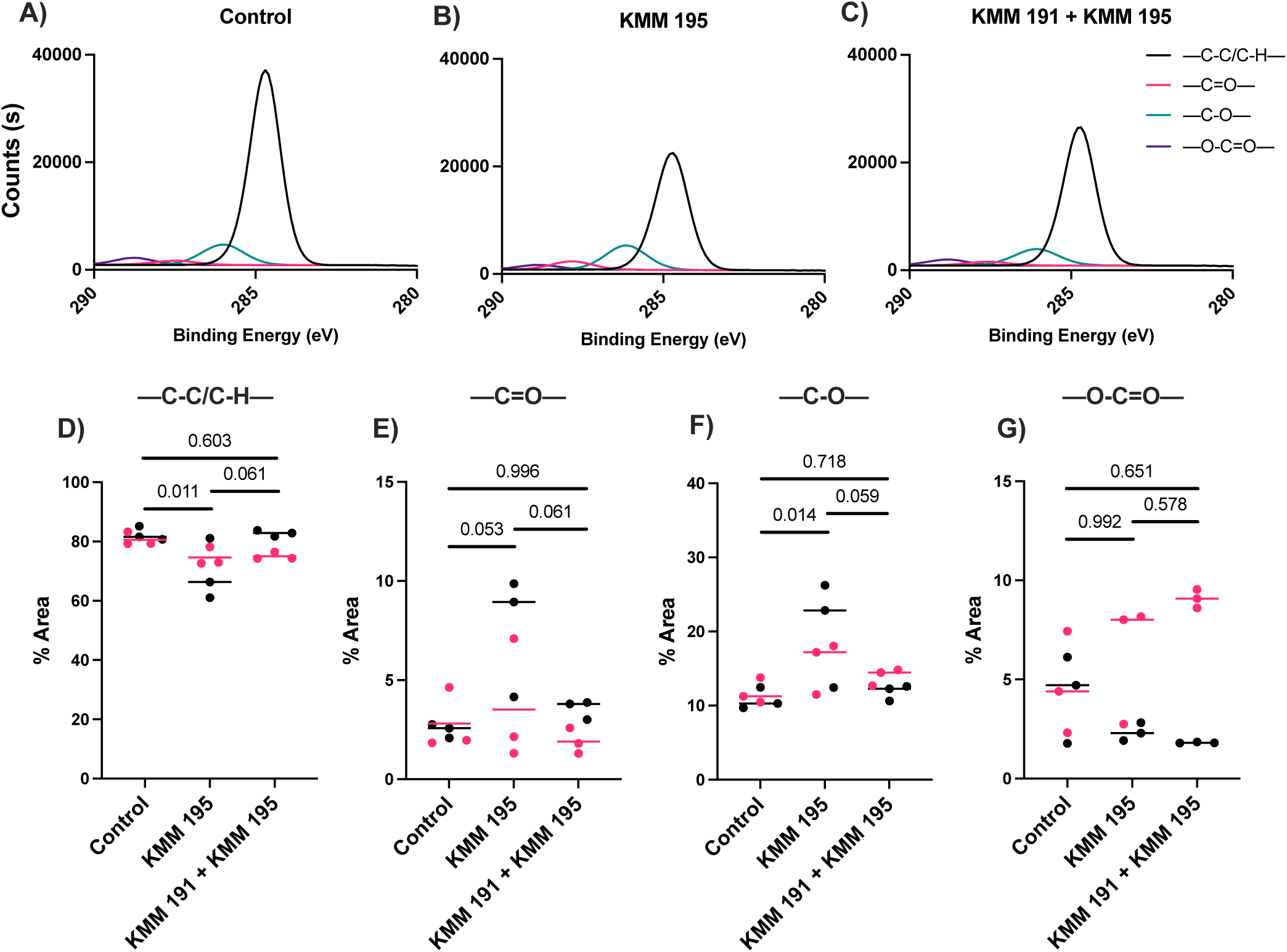
XPS analysis shows that KMM 195 induces oxidative surface modification of PP, while this effect is reduced in co-culture with KMM 191. **A-C)** High resolution XPS C 1s of PP films from the control, KMM 195, and the KMM 191 + KMM 195 co-culture **D-G)** Relative peak area percentage of C-C/C-H, C=O, C-O, and O-C=O groups from C 1s peak deconvolution. Each point represents a technical replicate (triplicate XPS scans), with colors distinguishing biological replicates. Lines represent the median for each biological replicate.

We used PERMANOVA to test for overall differences in multivariate comparisons to determine whether the entire surface chemistry profile shifts across treatments. Pairwise comparisons revealed that the KMM 195 treatment group differed significantly from the control (F = 4.71, R^2^ = 0.320, p = 0.039). In contrast, no significant differences were detected between the control and the KMM 191 + KMM 195 treatment group (F = 0.426, R^2^ = 0.041, p = 0.604), suggesting that co-culture conditions mitigate these changes in the surface chemistry. When directly comparing KMM 195 to the KMM 191 + KMM 195 treatment group, the difference was not statistically significant (p = 0.127) but exhibited a moderate effect size (F = 2.54, R^2^ = 0.203), still suggesting the chemical changes observed in monoculture are attenuated when KMM 195 is co-cultured with KMM 191.

To determine whether these chemical modifications were reflected into observable differences on the plastic surface, scanning electron microscopy (SEM) was performed on the same PP films. SEM revealed increased surface roughness and structural disruption in the cross-section of the films incubated with KMM 195 (Figure 9B) compared to the intact surface of the uninoculated control (Figure 9A). In contrast, the PP film incubated with the KMM 191 and KMM 195 co-culture revealed decreased surface roughness with a similar surface texture to the control (Figure 9C), supporting the hypothesis that KMM 191 inhibits plastic degradation.

**Figure 9.**
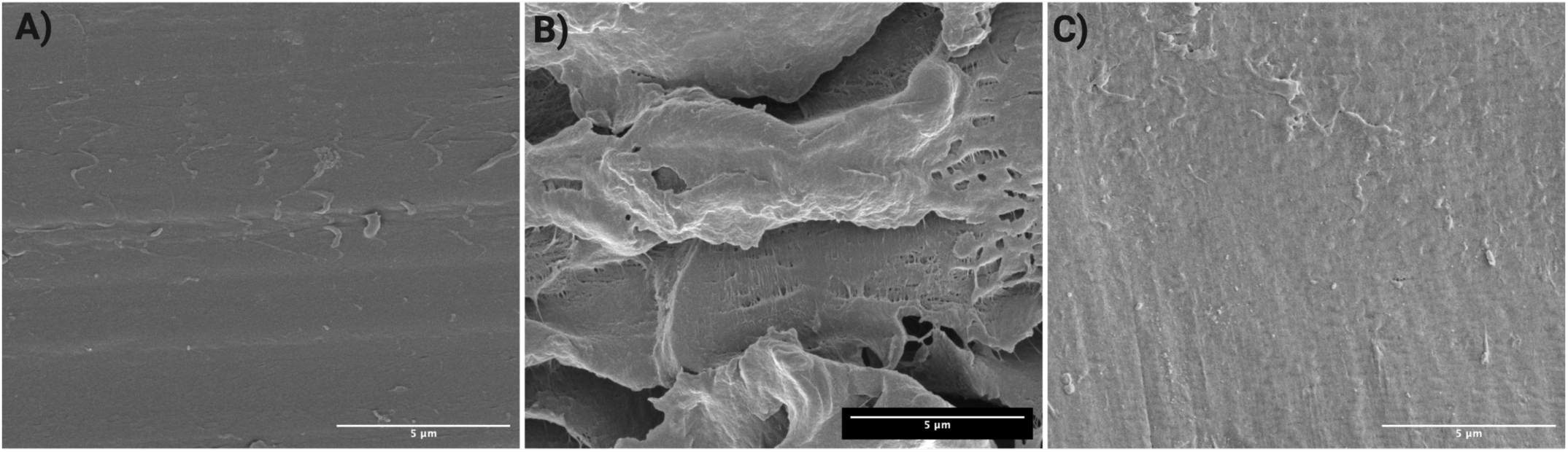
SEM images of PP film cross-sections. **A)** Uninoculated control showing a smooth and intact surface. **B)** PP film incubated with KMM 195 displaying increased surface roughness after 8 weeks of incubation in MSM. **C)** PP film incubated with KMM 191 and KMM 195 co-culture displaying decreased surface roughness after 8 weeks of incubation in MSM.

Overall, these morphological changes are consistent with the chemical changes shown through ATR-FTIR and XPS, further indicating that incubation with KMM 195 promotes PP oxidation and degradation under monoculture conditions. These surface-level observations parallel the enhanced lipase activity and biofilm formation observed for KMM 195 in monoculture, and the suppression of these activities in co-culture, linking enzymatic function and surface colonization dynamics to the resulting degradation phenotype.

## Discussion

In this study, we explored the enzymatic activity, biofilm formation, and active degradation potential of 184 bacterial isolates from the guts of threespine stickleback collected across six cold-water locations in Alaska. Our findings demonstrate that while some isolates exhibit biofilm formation and hydrolytic enzyme activity, these traits did not universally translate to detectable plastic degradation. Importantly, one isolate (KMM 195, *P. trivialis*) exhibited clear evidence of polypropylene surface modifications, indicating active plastic degradation. While many *Pseudomonas* species have been documented to degrade plastics^15,21,60,61^, *P. trivialis* has not been previously associated with this activity and has only been described from the phyllosphere of grasses^62^, highlighting a novel capability in this species and a unique strain from a cold-water environment where no plastic degraders have been identified. Other organisms have been shown to have a microbiome that degrades plastics; for example, multiple types of insect larvae have gut microbes capable of degrading plastics such as polyethylene and polystyrene^52,63,64^. Together, these findings suggest while plastic degradation is not universal, individual strains with enhanced capabilities may contribute to the response of the gut microbiome to plastic exposure. This activity may also provide a protective role for the host.

Strains such as KMM 195 could metabolize polymer byproducts that accumulate and localize in the gut, thereby reducing exposure to these compounds that would otherwise accumulate and impair host physiology. Because host-associated microbes are known to minimize effects of xenobiotics and dietary stressors^65,66^, plastic degradation within the gut would represent a novel extension of this protective role in cold-water environments.

Beyond monoculture experiments, our co-culture assays revealed that microbial interactions can significantly influence plastic degradation potential. Co-cultures with KMM 195 exhibited suppressed plastic degradation potential, notably in lipase activity and biofilm formation on PP when co-cultured with KMM 191 and KMM 230 (*P. germanica)*. Because these three strains were isolated from the same stickleback population, these results emphasize that members of the same microbiota could directly suppress the degradation potential of one another, and KMM 195 may possess intrinsic plastic-degrading capabilities. It is also possible that the introduction of another microbial species could counteract this suppression, either by inhibiting the inhibitory strain or by restoring degradative activity through cross-feeding or signaling interactions^67,68^. Antagonistic microbial interactions could constrain the expression of degradative enzymes, potentially through resource competition, quorum sensing interference or secretion of inhibitory enzymes, all of which have been observed in previous studies^69–72^. This finding emphasizes the need to evaluate the effects of microbial consortia rather than monocultures in environmental settings when exploring plastic degradation processes as degradation rates is likely shaped by complex networks of cooperation and inhibition^73,74^. This consideration is particularly important in cold-water settings where lower temperatures would likely slow chemical weathering and enzymatic reactions^75–77^, contributing to prolonged persistence of environmental plastics.

The identification of a cold-water *P. trivialis* strain with plastic-degrading potential brings potential for biotechnological applications. This observation expands the metabolic versatility of the species originally isolated in the phyllosphere in a warmer climate^62^ and suggests that this cold-adapted species could represent a group of enzymes with plastic degrading potential in low-temperature environments. Psychrophilic enzymes with plastic-degrading activity could offer advantages for bioremediation under ambient temperatures where thermophilic enzymes are less effective. Moreover, advances in enzyme engineering have demonstrated the potential of increasing hydrolase activity for polymer degradation^24,30–32,78^, highlighting the potential for psychrophilic enzymes as candidates for biotechnological advances.

While our assays provided low-cost, scalable methods to assess plastic degradation potential through enzymatic assays and biofilm formation on plastic films, they do not provide information on the specific enzymes or metabolic pathways responsible for degradation.

Although many hydrolases have broad substrate specificity, few hydrolases have been linked to specific plastic-degrading functions^18,33^. Future studies will combine whole genome sequencing, transcriptomics, and targeted enzyme assays to identify potential degradation pathways^79^.

Moreover, the plastic films used in this study were produced under controlled conditions that lack the additives, dyes, and weathering effects that characterize most environmental plastics^2,77,80,81^. Testing gut isolates on weathered plastics with environmental modifications would determine whether the degradation activities observed here reflect ecologically relevant degradation processes. Additional approaches such as stable isotope probing with ^13^C-labeled plastics could confirm the uptake of degradation products into the microbe^82^. Mock community experiments that combine isolates with complementary traits^12,14,15,22,83^ would also help determine whether cooperative interactions could minimize the inhibitory interactions observed in co-culture experiments. Ultimately, these directions will clarify whether this degradation potential could be translated into a complex natural ecosystem. By addressing this, future work will help determine whether the gut microbiome of cold-water fish represents a significant source of novel enzymes or communities for bioremediation purposes.

## Conclusion

Overall, our results indicate that the threespine stickleback gut microbiome contains bacterial strains with plastic degrading potential and could play a protective role by metabolizing polymer byproducts in cold-water environments, offering a potential avenue for bioremediation. These findings highlight the potential and complexity of microbiome-driven bioremediation.

Individual strains such as KMM 195 can degrade plastic, however plastic degrading activity is shaped by intra-community interactions that may either enhance or suppress this capability.

Together, these results contribute to a growing understanding of how host-associated microbes may influence interactions with environmental plastics and that plastic degradation is driven by microbial community interactions and point toward the gut microbiome as a potential source for harboring novel cold-adapted plastic degrading bacteria.

