## Supplemental Tables 1 & 2 for "Cold-water gut isolate from threespine stickleback *(Gasterosteus aculeatus)* reveals polypropylene surface oxidation and co-culture inhibition"

| Supplementary Table 1 \| Identifications of all 183 isolates exhibiting lipase activity, along with mean hydrolysis area from esterase assays, biofilm formation measurements, and carbonyl index values from FTIR. Blanks indicate that the corresponding test was not performed for that isolate. | | | | | | | | | | | | |
| --- | --- | --- | --- | --- | --- | --- | --- | --- | --- | --- | --- | --- |
| Microbe ID | **Population** | **Species** | **Lipase Activity** | **1% Esterase Hydrolysis Area** | **2% Esterase Hydrolysis Area** | **No Plastic RBF** | **PET Well RBF** | **PET Film RBF** | **PP Well RBF** | **PP Film RBF** | **PET Carbonyl Index** | **PP Carbonyl Index** |
| KMM 183 | WC | *Erwinia billingiae* | - | 0.0 ± 0.0 | 6.8 ± 5.9 |  |  |  |  |  | 2.528 ± 0.297 | 0.026 ± 0.028 |
| KMM 184 | WC | *Deefgea rivuli* | - | 0.0 ± 0.0 | 0.0 ± 0.0 |  |  |  |  |  |  |  |
| KMM 185 | WC | *Pseudomonas helleri* | - | 40.3 ± 8.7 | 45.3 ± 8.7 | 0.089 ± 0.034 | 0.083 ± 0.070 | 0.089 ± 0.081 | 0.107 ± 0.063 | 0.145 ± 0.042 | 2.791 ± 0.338 | 0.002 ± 0.021 |
| KMM 186 | WC | *Serratia sp.* | - | 72.8 ± 10.4 | 72.8 ± 10.4 |  |  |  |  |  |  |  |
| KMM 187 | WC | *Bacillus mycoides* | - | 55.8 ± 9.5 | 72.8 ± 10.4 |  |  |  |  |  |  |  |
| KMM 188 | WC | *Pantoea sp.* | - | 27.5 ± 20.6 | 3.4 ± 5.9 |  |  |  |  |  |  |  |
| KMM 189 | WC | *Shewanella putrefaciens* | - | 91.9 ± 21.8 | 97.9 ± 11.4 |  |  |  |  |  |  |  |
| KMM 191 | WC | *Pseudomonas germanica* | + | 299.8 ± 81.4 | 299.3 ± 76.2 | 0.076 ± 0.019 | 0.074 ± 0.009 | 0.043 ± 0.040 | 0.074 ± 0.010 | 0.036 ± 0.041 | 2.697 ± 0.198 | -0.003 ± 0.021 |
| KMM 192 | WC | *Serratia proteamaculans* | - | 14.1 ± 6.8 | 0.0 ± 0.0 |  |  |  |  |  |  |  |
| KMM 193 | WC | *Pseudomonas fragi* | - | 40.3 ± 8.7 | 40.3 ± 8.7 | 0.040 ± 0.028 | 0.036 ± 0.023 | 0.077 ± 0.060 | 0.053 ± 0.037 | 0.015 ± 0.011 | 2.717 ± 0.334 | 0.005 ± 0.018 |
| KMM 194 | WC | *Curtobacterium or Pantoea* | - | 10.7 ± 11.0 | 3.4 ± 5.9 |  |  |  |  |  |  |  |
| KMM 195 | WC | *Pseudomonas trivialis* | + | 286.4 ± 31.4 | 285.9 ± 0.0 | 0.047 ± 0.022 | 0.049 ± 0.016 | 0.073 ± 0.080 | 0.051 ± 0.021 | 0.043 ± 0.042 | 2.802 ± 0.117 | 0.059 ± 0.071 |
| KMM 196 | WC | *Shewanella vesiculosa* | - | 0.0 ± 0.0 | 0.0 ± 0.0 |  |  |  |  |  | 2.653 ± 0.297 | 0.010 ± 0.033 |
| KMM 197 | WC | *Aeromonas salmonicida* | + | 149.0 ± 23.6 | 173.3 ± 25.2 | 0.151 ± 0.098 | 0.091 ± 0.029 | 0.084 ± 0.054 | 0.095 ± 0.029 | 0.030 ± 0.028 | 2.956 ± 0.716 | 0.014 ± 0.037 |
| KMM 198 | WC | *Pantoea sp.* | - | 50.8 ± 15.8 | 86.9 ± 37.7 |  |  |  |  |  | 2.546 ± 0.215 | 0.026 ± 0.026 |
| KMM 199 | WC | *Serratia aquatilis* | - | 73.9 ± 27.8 | 85.4 ± 18.9 |  |  |  |  |  | 2.531 ± 0.259 | 0.009 ± 0.026 |
| KMM 200 | WC | *Erwinia persicina* | - | 26.4 ± 7.7 | 35.9 ± 14.2 | 0.114 ± 0.106 | 0.021 ± 0.008 | 0.088 ± 0.031 | 0.031 ± 0.004 | 0.048 ± 0.031 |  |  |
| KMM 202 | WC | *Pantoea agglomerans* | - | 55.8 ± 9.5 | 26.4 ± 7.7 | 0.124 ± 0.013 | 0.130 ± 0.022 | 0.062 ± 0.042 | 0.133 ± 0.028 | 0.080 ± 0.082 | 2.665 ± 0.350 | 0.000 ± 0.021 |
| KMM 203 | WC | *Arthrobacter luteolus* | - | 30.9 ± 7.7 | 23.5 ± 20.4 | 0.005 ± 0.004 | 0.012 ± 0.006 | -0.007 ± 0.004 | 0.008 ± 0.002 | 0.005 ± 0.006 |  |  |
| KMM 204 | WC | *Pseudomonas azotoformans* | + | 119.1 ± 25.3 | 118.6 ± 12.2 | 0.069 ± 0.022 | 0.083 ± 0.034 | 0.113 ± 0.101 | 0.089 ± 0.033 | 0.169 ± 0.204 | 2.831 ± 0.136 | -0.013 ± 0.027 |
| KMM 205 | WC | *Brochothrix thermosphacta* | - | 55.8 ± 9.5 | 50.3 ± 0.0 | 0.124 ± 0.016 | 0.128 ± 0.015 | 0.055 ± 0.033 | 0.146 ± 0.016 | 0.037 ± 0.019 |  |  |
| KMM 206 | WC | *Buttiauxella sp.* | - | 7.3 ± 12.7 | 26.4 ± 7.7 | 0.124 ± 0.002 | 0.121 ± 0.019 | 0.033 ± 0.012 | 0.114 ± 0.005 | 0.028 ± 0.005 |  |  |
| KMM 207 | WC | *Pantoea agglomerans* | - | 16.8 ± 29.0 | 35.3 ± 0.0 | 0.132 ± 0.012 | 0.159 ± 0.024 | 0.100 ± 0.047 | 0.130 ± 0.013 | 0.018 ± 0.019 |  |  |
| KMM 208 | WC | *Paenibacillus xylanexedens* | - | 78.8 ± 10.4 | 50.3 ± 0.0 | 0.038 ± 0.006 | 0.021 ± 0.002 | 0.002 ± 0.009 | 0.029 ± 0.008 | -0.002 ± 0.000 |  |  |
| KMM 209 | WC | *Erwinia billingiae* | - | 14.7 ± 12.7 | 22.0 ± 0.0 |  |  |  |  |  |  |  |
| KMM 210 | BL | *Erwinia billingiae* | - | 19.1 ± 17.8 | 30.9 ± 7.7 |  |  |  |  |  |  |  |
| KMM 211 | BL | *Deefgea piscis* | - | 93.5 ± 38.9 | 62.3 ± 25.1 |  |  |  |  |  |  |  |
| KMM 212 | BL | *Pseudomonas paraversuta* | - | 127.2 ± 36.7 | 81.4 ± 40.0 | 0.101 ± 0.019 | 0.173 ± 0.011 | 0.011 ± 0.010 | 0.141 ± 0.016 | 0.012 ± 0.018 |  |  |
| KMM 213 | Mud | *Koukoulia aurantiaca* | - | 0.0 ± 0.0 | 0.0 ± 0.0 |  |  |  |  |  |  |  |
| KMM 214 | Mud | *Aeromonas sobria* | + | 215.2 ± 97.1 | 74.9 ± 35.7 | 0.060 ± 0.019 | 0.058 ± 0.020 | 0.020 ± 0.015 | 0.058 ± 0.021 | 0.012 ± 0.014 | 2.851 ± 0.165 | 0.003 ± 0.027 |
| KMM 216 | BL | *Deefgea piscis* | - | 0.0 ± 0.0 | 0.0 ± 0.0 |  |  |  |  |  |  |  |
| KMM 217 | BL | *Aeromonas salmonicida* | + | 181.4 ± 15.0 | 152.1 ± 65.0 | 0.064 ± 0.025 | 0.061 ± 0.012 | 0.018 ± 0.016 | 0.061 ± 0.013 | 0.087 ± 0.217 | 2.783 ± 0.122 | -0.015 ± 0.021 |
| KMM 218 | Mud | *Aeromonas sobria* | + | 226.2 ± 0.0 | 150.0 ± 42.1 | 0.038 ± 0.016 | 0.033 ± 0.014 | 0.012 ± 0.013 | 0.032 ± 0.011 | 0.007 ± 0.016 |  |  |
| KMM 219 | BL | *Shewanella oneidensis* | + | 63.1 ± 55.5 | 50.8 ± 15.8 | 0.007 ± 0.004 | 0.018 ± 0.017 | 0.037 ± 0.046 | 0.018 ± 0.009 | 0.042 ± 0.051 | 2.877 ± 0.192 | -0.025 ± 0.012 |
| KMM 220 | BL | *Deefgea piscis* | - | 0.0 ± 0.0 | 0.0 ± 0.0 |  |  |  |  |  |  |  |
| KMM 221 | BL | *Providencia alcalifaciens* | - | 0.0 ± 0.0 | 3.4 ± 5.9 |  |  |  |  |  |  |  |
| KMM 222 | Mud | *Pseudomonas peli* | - | 28.3 ± 49.0 | 78.8 ± 10.4 | 0.052 ± 0.027 | 0.065 ± 0.016 | 0.041 ± 0.042 | 0.056 ± 0.006 | 0.122 ± 0.091 |  |  |
| KMM 223 | BL | *Deefgea piscis* | - | 0.0 ± 0.0 | 0.0 ± 0.0 |  |  |  |  |  |  |  |
| KMM 224 | BL | *Deefgea piscis* | - | 0.0 ± 0.0 | 0.0 ± 0.0 |  |  |  |  |  |  |  |
| KMM 225 | Mud | *Aeromonas veronii* | + | 161.5 ± 146.5 | 141.4 ± 27.2 |  |  |  |  |  |  |  |
| KMM 226 | WC | *Serratia oryzae* | - | 50.5 ± 44.7 | 68.4 ± 31.3 |  |  |  |  |  |  |  |
| KMM 227 | WC | *Bacillus mycoides* | - | 61.3 ± 9.5 | 45.3 ± 8.7 | 0.001 ± 0.003 | 0.001 ± 0.005 | 0.080 ± 0.076 | 0.012 ± 0.002 | 0.065 ± 0.106 |  |  |
| KMM 228 | WC | *Pantoea agglomerans* | - | 18.1 ± 6.8 | 30.9 ± 7.7 |  |  |  |  |  |  |  |
| KMM 229 | WC | *Shewanella putrefaciens* | - | 50.5 ± 44.7 | 62.3 ± 25.1 |  |  |  |  |  |  |  |
| KMM 230 | WC | *Pseudomonas germanica* | + | 227.8 ± 47.6 | 156.5 ± 14.1 | 0.076 ± 0.019 | 0.066 ± 0.014 | 0.025 ± 0.018 | 0.075 ± 0.019 | 0.028 ± 0.031 | 2.796 ± 0.216 | 0.012 ± 0.023 |
| KMM 231 | WC | *Serratia quinivorans* | - | 6.8 ± 5.9 | 6.8 ± 5.9 |  |  |  |  |  |  |  |
| KMM 232 | WC | *Pseudomonas bubulae* | - | 18.1 ± 6.8 | 10.2 ± 0.0 | 0.025 ± 0.014 | 0.024 ± 0.019 | 0.068 ± 0.055 | 0.025 ± 0.014 | 0.030 ± 0.033 | 2.843 ± 0.097 | -0.019 ± 0.035 |
| KMM 233 | WC | *Pantoea agglomerans* | - | 10.2 ± 0.0 | 3.4 ± 5.9 |  |  |  |  |  |  |  |
| KMM 234 | WC | *Shewanella vesiculosa* | - | 0.0 ± 0.0 | 0.0 ± 0.0 |  |  |  |  |  |  |  |
| KMM 235 | WC | *Aeromonas salmonicida* | + | 84.8 ± 0.0 | 73.3 ± 19.9 |  |  |  |  |  |  |  |
| KMM 236 | WC | *Pantoea agglomerans* | - | 22.0 ± 0.0 | 22.0 ± 0.0 | 0.011 ± 0.009 | 0.012 ± 0.006 | 0.011 ± 0.014 | 0.026 ± 0.035 | 0.008 ± 0.017 | 2.855 ± 0.156 | -0.023 ± 0.014 |
| KMM0237 | WC | *Serratia aquatilis* | - | 40.3 ± 8.7 | 26.4 ± 7.7 |  |  |  |  |  |  |  |
| KMM0238 | WC | *Erwinia persicina* | - | 3.4 ± 5.9 | 0.0 ± 0.0 |  |  |  |  |  | 2.763 ± 0.114 | 0.001 ± 0.025 |
| KMM0239 | WC | *Pantoea agglomerans* | - | 22.5 ± 12.6 | 10.2 ± 0.0 |  |  |  |  |  |  |  |
| KMM0240 | WC | *Pantoea agglomerans* | - | 14.1 ± 6.8 | 14.1 ± 6.8 |  |  |  |  |  |  |  |
| KMM0241 | WC | *Arthrobacter luteolus* | - | 0.0 ± 0.0 | 0.0 ± 0.0 |  |  |  |  |  |  |  |
| KMM0243 | WC | *Brochothrix thermosphacta* | - | 26.4 ± 7.7 | 14.1 ± 6.8 |  |  |  |  |  |  |  |
| KMM0244 | WC | *Buttiauxella ferragutiae* | - | 10.2 ± 0.0 | 0.0 ± 0.0 | 0.249 ± 0.034 | 0.128 ± 0.016 | 0.024 ± 0.027 | 0.127 ± 0.015 | 0.025 ± 0.021 |  |  |
| KMM0245 | WC | *Pantoea agglomerans* | - | 18.1 ± 6.8 | 10.2 ± 0.0 | 0.119 ± 0.022 | 0.149 ± 0.019 | 0.038 ± 0.010 | 0.136 ± 0.007 | 0.003 ± 0.005 | 2.930 ± 0.128 | -0.009 ± 0.011 |
| KMM0246 | WC | *Erwinia billingiae* | - | 0.0 ± 0.0 | 0.0 ± 0.0 |  |  |  |  |  |  |  |
| KMM0247 | WC | *Pantoea agglomerans* | - | 6.8 ± 5.9 | 6.8 ± 5.9 |  |  |  |  |  |  |  |
| KMM0248 | WC | *Flavobacterium hydatis* | + | 61.3 ± 9.5 | 55.8 ± 9.5 |  |  |  |  |  |  |  |
| KMM0249 | Mud | *Pseudomonas sesami* | + | 266.5 ± 47.0 | 191.1 ± 39.5 | 0.114 ± 0.019 | 0.109 ± 0.016 | 0.028 ± 0.008 | 0.105 ± 0.019 | 0.088 ± 0.038 |  |  |
| KMM0250 |  | *Erwinia billingiae* | - | 0.0 ± 0.0 | 0.0 ± 0.0 |  |  |  |  |  |  |  |
| KMM0251 |  | *Erwinia billingiae* | - | 0.0 ± 0.0 | 0.0 ± 0.0 |  |  |  |  |  |  |  |
| KMM0252 |  | *Erwinia billingiae* | - | 0.0 ± 0.0 | 0.0 ± 0.0 |  |  |  |  |  |  |  |
| KMM0254 |  | *Erwinia billingiae* | - | 0.0 ± 0.0 | 0.0 ± 0.0 |  |  |  |  |  |  |  |
| KMM0255 | CL | *Lysinibacillus sphaericus* | + | 97.9 ± 11.4 | 66.8 ± 0.0 | 0.158 ± 0.103 | 0.032 ± 0.005 | -0.002 ± 0.002 | 0.025 ± 0.006 | 0.051 ± 0.103 |  |  |
| KMM0257 | CL | *Aeromonas salmonicida* | - | 97.9 ± 11.4 | 91.9 ± 21.8 |  |  |  |  |  |  |  |
| KMM0258 | CL | *Sporosarcina aquimarina* | - | 0.0 ± 0.0 | 0.0 ± 0.0 |  |  |  |  |  | 2.789 ± 0.103 | -0.001 ± 0.017 |
| KMM0259 | CL | *Sporosarcina aquimarina* | - | 10.7 ± 11.0 | 18.1 ± 6.8 |  |  |  |  |  |  |  |
| KMM0260 | CL | *Aeromonas popoffii* | - | 50.3 ± 0.0 | 55.8 ± 9.5 |  |  |  |  |  |  |  |
| KMM0262 | CL | *Pseudomonas azotoformans* | + | 157.6 ± 37.4 | 156.5 ± 14.1 |  |  |  |  |  |  |  |
| KMM0265 | CL | *Aeromonas salmonicida* | - | 91.9 ± 21.8 | 78.8 ± 10.4 |  |  |  |  |  | 2.899 ± 0.126 | -0.002 ± 0.017 |
| KMM0266 | CL | *Solibacillus silvestris* | + | 14.1 ± 6.8 | 22.0 ± 0.0 |  |  |  |  |  |  |  |
| KMM0267 | CL | *Pseudomonas azotoformans* | + | 61.8 ± 19.9 | 72.8 ± 10.4 | 0.055 ± 0.007 | 0.078 ± 0.036 | 0.226 ± 0.087 | 0.091 ± 0.051 | 0.063 ± 0.055 | 2.907 ± 0.152 | -0.023 ± 0.020 |
| KMM0270 | CL | *Pseudomonas azotoformans* | + | 173.3 ± 25.2 | 118.6 ± 12.2 |  |  |  |  |  |  |  |
| KMM0271 | CL | *Oerskovia turbata* | - | 10.2 ± 0.0 | 10.2 ± 0.0 |  |  |  |  |  |  |  |
| KMM0272 | CL | *Bacillus cereus* | - | 55.8 ± 9.5 | 55.8 ± 9.5 |  |  |  |  |  |  |  |
| KMM0273 | CL | *Aeromonas allosaccharophila* | + | 111.6 ± 12.2 | 104.5 ± 0.0 | 0.054 ± 0.026 | 0.073 ± 0.022 | 0.034 ± 0.018 | 0.078 ± 0.027 | 0.023 ± 0.020 | 2.850 ± 0.146 | -0.039 ± 0.022 |
| KMM0274 | CL | *Sporosarcina aquimarina* | - | 15.2 ± 18.2 | 22.0 ± 0.0 |  |  |  |  |  |  |  |
| KMM0275 | CL | *Bacillus mycoides* | + | 112.6 ± 32.6 | 78.8 ± 10.4 | 0.015 ± 0.004 | 0.013 ± 0.004 | -0.003 ± 0.006 | 0.029 ± 0.017 | -0.017 ± 0.002 | 2.839 ± 0.203 | 0.010 ± 0.020 |
| KMM0276 | CL | *Sporosarcina aquimarina* | - | 35.9 ± 14.2 | 35.9 ± 14.2 | 0.081 ± 0.025 | 0.130 ± 0.020 | 0.017 ± 0.008 | 0.091 ± 0.014 | 0.030 ± 0.017 |  |  |
| KMM0277 | CL | *Bacillus mycoides* | + | 53.7 ± 71.7 | 91.4 ± 11.4 |  |  |  |  |  | 2.655 ± 0.445 | 0.021 ± 0.022 |
| KMM0279 | CL | *Solibacillus sp.* | + | 0.0 ± 0.0 | 0.0 ± 0.0 |  |  |  |  |  |  |  |
| KMM0280 | CL | *Solibacillus sp.* | + | 0.0 ± 0.0 | 3.4 ± 5.9 |  |  |  |  |  |  |  |
| KMM0281 | CL | *Solibacillus sp.* | + | 0.0 ± 0.0 | 0.0 ± 0.0 |  |  |  |  |  | 2.940 ± 0.109 | -0.028 ± 0.034 |
| KMM0282 | CL | *Sporosarcina aquimarina* | - | 0.0 ± 0.0 | 3.4 ± 5.9 |  |  |  |  |  |  |  |
| KMM0283 | CL | *Sporosarcina aquimarina* | - | 18.1 ± 6.8 | 14.1 ± 6.8 |  |  |  |  |  |  |  |
| KMM0284 | CL | *Aeromonas salmonicida* | - | 105.0 ± 20.5 | 78.8 ± 10.4 | 0.099 ± 0.024 | 0.100 ± 0.018 | 0.063 ± 0.043 | 0.096 ± 0.014 | 0.051 ± 0.073 | 2.852 ± 0.098 | -0.019 ± 0.028 |
| KMM0285 | CL | *Deefgea piscis* | + | 0.0 ± 0.0 | 0.0 ± 0.0 |  |  |  |  |  |  |  |
| KMM0286 | CL | *Aeromonas salmonicida* | - | 105.0 ± 20.5 | 91.4 ± 11.4 | 0.079 ± 0.023 | 0.105 ± 0.028 | 0.031 ± 0.020 | 0.100 ± 0.024 | 0.036 ± 0.058 | 2.982 ± 0.223 | -0.014 ± 0.025 |
| KMM0287 | CL | *Neissariaceae bacterium* | - | 14.1 ± 6.8 | 10.2 ± 0.0 |  |  |  |  |  |  |  |
| KMM0288 | CL | *Erwinia billingiae* | - | 72.8 ± 10.4 | 35.3 ± 0.0 |  |  |  |  |  |  |  |
| KMM0289 | CL | *Aeromonas veronii bv. veronii* | + | 140.8 ± 13.1 | 148.4 ± 0.0 | 0.096 ± 0.039 | 0.085 ± 0.019 | 0.038 ± 0.027 | 0.091 ± 0.025 | 0.035 ± 0.026 | 2.733 ± 0.463 | 0.017 ± 0.031 |
| KMM0290 | CL | *escherichia marmotae* | - | 0.0 ± 0.0 | 0.0 ± 0.0 |  |  |  |  |  |  |  |
| KMM0291 | CL | *chryseobacterium soli* | - | 26.4 ± 7.7 | 18.1 ± 6.8 |  |  |  |  |  |  |  |
| KMM0292 | CL | *Epilithonimonas ginsengisoli* | - | 26.4 ± 7.7 | 22.5 ± 12.6 |  |  |  |  |  | 2.806 ± 0.415 | 0.009 ± 0.025 |
| KMM0293 | CL | *Chromobacterium aquaticum* | + | 173.3 ± 25.2 | 182.5 ± 39.8 | 0.081 ± 0.050 | 0.060 ± 0.007 | 0.119 ± 0.107 | 0.065 ± 0.014 | 0.062 ± 0.069 | 2.650 ± 0.235 | 0.012 ± 0.015 |
| KMM0294 | CL | *Naganishia sp.* | - | 35.9 ± 14.2 | 10.7 ± 11.0 |  |  |  |  |  |  |  |
| KMM0295 | CL | *Pseudomonas alcaligenes* | - | 50.3 ± 0.0 | 35.3 ± 0.0 |  |  |  |  |  |  |  |
| KMM0296 | CL | *Rhodococcus erythropolis* | + | 61.3 ± 9.5 | 61.3 ± 9.5 |  |  |  |  |  |  |  |
| KMM0297 | CL | *Escherichia marmotae* | - | 0.0 ± 0.0 | 0.0 ± 0.0 |  |  |  |  |  |  |  |
| KMM0298 | CL | *Brochothrix thermosphacta* | - | 0.0 ± 0.0 | 0.0 ± 0.0 |  |  |  |  |  |  |  |
| KMM0299 | CL | *Rheinheimera mesophila* | - | 0.0 ± 0.0 | 0.0 ± 0.0 |  |  |  |  |  |  |  |
| KMM0300 | CL | *Pseudomonas abietaniphila* | - | 35.3 ± 0.0 | 35.3 ± 0.0 | 0.015 ± 0.008 | 0.004 ± 0.009 | 0.033 ± 0.021 | 0.029 ± 0.054 | 0.106 ± 0.166 |  |  |
| KMM0301 | CL | *Pseudomonas protegens* | + | 173.3 ± 25.2 | 165.7 ± 37.0 | 0.059 ± 0.025 | 0.049 ± 0.017 | 0.055 ± 0.027 | 0.045 ± 0.019 | 0.044 ± 0.018 | 2.913 ± 0.161 | 0.000 ± 0.037 |
| KMM0302 | CL | *Escherichia marmotae* | - | 0.0 ± 0.0 | 0.0 ± 0.0 |  |  |  |  |  |  |  |
| KMM352 | RS | *Shewanella putrefaciens* | - | 18.1 ± 6.8 | 31.9 ± 20.3 |  |  |  |  |  | 2.903 ± 0.261 | 0.017 ± 0.015 |
| KMM353 | RS | *Aeromonas sobria* | + | 118.6 ± 12.2 | 133.8 ± 25.3 |  |  |  |  |  |  |  |
| KMM354 | RS | *Shewanella putrefaciens* | + | 35.9 ± 14.2 | 26.4 ± 7.7 |  |  |  |  |  | 2.981 ± 0.208 | 0.015 ± 0.025 |
| KMM355 | RS | *Aeromonas sobria* | + | 126.2 ± 22.0 | 105.0 ± 20.5 |  |  |  |  |  | 2.840 ± 0.146 | 0.004 ± 0.016 |
| KMM356 | RS | *Shewanella putrefaciens* | + | 26.4 ± 7.7 | 26.4 ± 7.7 | 0.061 ± 0.003 | 0.073 ± 0.013 | 0.165 ± 0.057 | 0.089 ± 0.008 | 0.081 ± 0.010 | 2.812 ± 0.161 | 0.010 ± 0.014 |
| KMM357 | RS | *Aeromonas sobria* | + | 111.6 ± 12.2 | 92.4 ± 30.2 | 0.027 ± 0.002 | 0.024 ± 0.001 | 0.021 ± 0.015 | 0.035 ± 0.035 | -0.001 ± 0.009 | 2.975 ± 0.177 | 0.000 ± 0.018 |
| KMM358 | RS | *Aeromonas veronii* | + | 118.6 ± 12.2 | 106.0 ± 36.7 | 0.104 ± 0.016 | 0.102 ± 0.007 | 0.095 ± 0.034 | 0.077 ± 0.002 | 0.042 ± 0.026 | 2.942 ± 0.191 | 0.005 ± 0.016 |
| KMM359 | RS | *Shewanella putrefaciens* | + | 30.9 ± 7.7 | 31.4 ± 16.3 |  |  |  |  |  |  |  |
| KMM360 | RS | *Serratia fonticola* | - | 91.4 ± 11.4 | 31.9 ± 20.3 |  |  |  |  |  |  |  |
| KMM361 | RS | *Serratia fonticola* | - | 84.8 ± 0.0 | 35.9 ± 14.2 |  |  |  |  |  |  |  |
| KMM362 | RS | *Serratia fonticola* | - | 84.8 ± 0.0 | 30.9 ± 7.7 |  |  |  |  |  |  |  |
| KMM363 | RS | *Serratia fonticola* | - | 91.9 ± 21.8 | 30.9 ± 7.7 |  |  |  |  |  |  |  |
| KMM364 | RS | *Aeromonas veronii* | + | 217.0 ± 15.9 | 148.4 ± 0.0 | 0.114 ± 0.047 | 0.088 ± 0.011 | 0.115 ± 0.093 | 0.111 ± 0.026 | 0.076 ± 0.032 | 2.991 ± 0.150 | 0.009 ± 0.016 |
| KMM365 | RS | *Aeromonas veronii* | + | 181.4 ± 15.0 | 97.9 ± 11.4 |  |  |  |  |  |  |  |
| KMM366 | RS | *Aeromonas veronii* | + | 199.2 ± 26.7 | 126.2 ± 22.0 |  |  |  |  |  |  |  |
| KMM367 | RS | *Aeromonas veronii* | + | 226.2 ± 0.0 | 125.7 ± 0.0 | 0.097 ± 0.027 | 0.097 ± 0.026 | 0.103 ± 0.036 | 0.099 ± 0.023 | 0.149 ± 0.183 | 3.096 ± 0.183 | 0.033 ± 0.048 |
| KMM368 | RS | *Serratia fonticola* | - | 104.5 ± 0.0 | 30.9 ± 7.7 | 0.057 ± 0.012 | 0.088 ± 0.024 | 0.053 ± 0.003 | 0.044 ± 0.003 | 0.089 ± 0.020 | 2.908 ± 0.234 | 0.016 ± 0.021 |
| KMM369 | RS | *Serratia fonticola* | - | 104.5 ± 0.0 | 30.9 ± 7.7 | 0.045 ± 0.014 | 0.046 ± 0.014 | 0.025 ± 0.021 | 0.049 ± 0.013 | 0.048 ± 0.037 |  |  |
| KMM370 | RS | *Serratia fonticola* | - | 35.3 ± 0.0 | 30.9 ± 7.7 |  |  |  |  |  |  |  |
| KMM371 | RS | *Serratia fonticola* | - | 30.9 ± 7.7 | 22.0 ± 0.0 |  |  |  |  |  |  |  |
| KMM372 | RS | *Aeromonas veronii* | + | 148.4 ± 0.0 | 148.4 ± 0.0 |  |  |  |  |  |  |  |
| KMM373 | RS | *Aeromonas veronii* | + | 148.4 ± 0.0 | 140.8 ± 13.1 |  |  |  |  |  |  |  |
| KMM374 | RS | *Aeromonas veronii* | + | 140.8 ± 13.1 | 140.8 ± 13.1 |  |  |  |  |  | 2.918 ± 0.228 | 0.006 ± 0.027 |
| KMM375 | RS | *Aeromonas veronii* | + | 148.4 ± 0.0 | 148.4 ± 0.0 |  |  |  |  |  | 2.803 ± 0.117 | 0.018 ± 0.023 |
| KMM376 | RS | *Aeromonas veronii* | + | 217.0 ± 15.9 | 111.6 ± 12.2 | 0.080 ± 0.016 | 0.082 ± 0.019 | 0.066 ± 0.035 | 0.086 ± 0.015 | 0.080 ± 0.042 |  |  |
| KMM377 | RS | *Aeromonas veronii* | + | 181.4 ± 15.0 | 111.6 ± 12.2 |  |  |  |  |  |  |  |
| KMM378 | RS | *Rossellomorea vietnamensis* | - | 0.0 ± 0.0 | 0.0 ± 0.0 |  |  |  |  |  |  |  |
| KMM379 | RS | *Aeromonas veronii* | + | 181.9 ± 29.0 | 111.6 ± 12.2 | 0.140 ± 0.009 | 0.172 ± 0.008 | 0.055 ± 0.009 | 0.161 ± 0.004 | 0.061 ± 0.010 |  |  |
| KMM380 | RS | *Aeromonas veronii* | + | 199.2 ± 26.7 | 133.3 ± 13.1 | 0.068 ± 0.010 | 0.075 ± 0.007 | 0.052 ± 0.011 | 0.071 ± 0.018 | 0.070 ± 0.029 |  |  |
| KMM381 | RS | *Shewanella putrefaciens* | - | 61.3 ± 9.5 | 30.9 ± 7.7 |  |  |  |  |  | 2.775 ± 0.109 | 0.006 ± 0.022 |
| KMM382 | RS | *Aeromonas veronii* | + | 148.4 ± 0.0 | 148.4 ± 0.0 |  |  |  |  |  |  |  |
| KMM383 | RS | *Aeromonas veronii* | + | 148.4 ± 0.0 | 148.4 ± 0.0 | 0.071 ± 0.003 | 0.072 ± 0.006 | 0.091 ± 0.043 | 0.071 ± 0.014 | 0.090 ± 0.009 |  |  |
| KMM384 | RS | *Shewanella putrefaciens* | + | 126.2 ± 22.0 | 30.9 ± 7.7 |  |  |  |  |  |  |  |
| KMM385 | RS | *Aeromonas veronii* | + | 236.9 ± 44.6 | 111.6 ± 12.2 | 0.155 ± 0.021 | 0.169 ± 0.021 | 0.040 ± 0.004 | 0.177 ± 0.013 | 0.052 ± 0.023 |  |  |
| KMM386 | RS | *Aeromonas veronii* | + | 276.7 ± 46.6 | 111.6 ± 12.2 | 0.060 ± 0.009 | 0.051 ± 0.002 | 0.082 ± 0.021 | 0.058 ± 0.004 | 0.051 ± 0.028 |  |  |
| KMM387 | RS | *Aeromonas veronii* | + | 276.7 ± 46.6 | 111.6 ± 12.2 | 0.070 ± 0.006 | 0.072 ± 0.011 | 0.078 ± 0.019 | 0.079 ± 0.007 | 0.062 ± 0.009 |  |  |
| KMM389 | RS | *Micrococcus luteus* | + | 0.0 ± 0.0 | 0.0 ± 0.0 | 0.014 ± 0.004 | 0.055 ± 0.018 | 0.004 ± 0.005 | 0.049 ± 0.032 | -0.003 ± 0.004 |  |  |
| KMM390 | RS | *Micrococcus luteus* | + | 67.3 ± 17.3 | 22.0 ± 0.0 | 0.071 ± 0.022 | 0.075 ± 0.040 | 0.000 ± 0.002 | 0.086 ± 0.052 | -0.005 ± 0.010 | 3.068 ± 0.126 | 0.023 ± 0.023 |
| KMM392 | BP | *Pantoea vagans* | - | 16.8 ± 29.0 | 18.1 ± 6.8 | 0.079 ± 0.008 | 0.072 ± 0.008 | 0.068 ± 0.020 | 0.069 ± 0.012 | 0.069 ± 0.013 |  |  |
| KMM393 | BP | *Pantoea vagans* | - | 22.3 ± 38.6 | 22.0 ± 0.0 | 0.098 ± 0.011 | 0.049 ± 0.002 | 0.044 ± 0.008 | 0.054 ± 0.015 | 0.064 ± 0.013 |  |  |
| KMM394 | BP | *Hafnia paralvei* | - | 11.8 ± 20.4 | 22.0 ± 0.0 |  |  |  |  |  |  |  |
| KMM395 | BP | *Hafnia paralvei* | - | 28.5 ± 25.8 | 14.1 ± 6.8 |  |  |  |  |  |  |  |
| KMM396 | BP | *Hafnia paralvei* | - | 14.7 ± 12.7 | 18.1 ± 6.8 |  |  |  |  |  |  |  |
| KMM397 | BP | *Pantoea vagans* | - | 23.5 ± 20.4 | 30.9 ± 7.7 | 0.054 ± 0.020 | 0.052 ± 0.005 | 0.035 ± 0.013 | 0.059 ± 0.013 | 0.089 ± 0.028 |  |  |
| KMM398 | BP | *Hafnia paralvei* | - | 55.8 ± 9.5 | 30.9 ± 7.7 |  |  |  |  |  |  |  |
| KMM399 | BP | *Serratia plymuthica* | - | 40.9 ± 16.3 | 26.4 ± 7.7 |  |  |  |  |  |  |  |
| KMM400 | BP | *Serratia plymuthica* | - | 39.0 ± 34.8 | 22.0 ± 0.0 |  |  |  |  |  |  |  |
| KMM401 | BP | *Serratia plymuthica* | + | 40.3 ± 8.7 | 22.0 ± 0.0 |  |  |  |  |  |  |  |
| KMM402 | BP | *Pantoea vagans* | - | 33.5 ± 29.0 | 26.4 ± 7.7 | 0.044 ± 0.006 | 0.055 ± 0.006 | 0.059 ± 0.001 | 0.033 ± 0.006 | 0.129 ± 0.048 | 2.877 ± 0.289 | 0.005 ± 0.017 |
| KMM403 | BP | *Pantoea vagans* | - | 40.0 ± 42.6 | 14.7 ± 12.7 |  |  |  |  |  |  |  |
| KMM404 | BP | *Pantoea vagans* | + | 50.3 ± 0.0 | 22.0 ± 0.0 |  |  |  |  |  |  |  |
| KMM405 | BP | *Micrococcus yunnanensis* | + | 105.0 ± 20.5 | 18.1 ± 6.8 |  |  |  |  |  |  |  |
| KMM406 | BP | *Paenibacillus dendritiformis* | + | 0.0 ± 0.0 | 0.0 ± 0.0 |  |  |  |  |  |  |  |
| KMM407 | BP | *Micrococcus antarcticus* | - | 0.0 ± 0.0 | 0.0 ± 0.0 |  |  |  |  |  |  |  |
| KMM408 | BP | *Micrococcus luteus* | - | 28.5 ± 25.8 | 22.0 ± 0.0 |  |  |  |  |  |  |  |
| KMM409 | BP | *Pantoea sp.* | + | 0.0 ± 0.0 | 0.0 ± 0.0 |  |  |  |  |  |  |  |
| KMM410 | BP | *Micrococcus antarcticus* | + | 3.4 ± 5.9 | 0.0 ± 0.0 |  |  |  |  |  |  |  |
| KMM411 | BP | *Paenibacillus odorifer* | - | 61.3 ± 9.5 | 40.3 ± 8.7 |  |  |  |  |  |  |  |
| KMM412 | BP | *Micrococcus* | - | 0.0 ± 0.0 | 0.0 ± 0.0 |  |  |  |  |  |  |  |
| KMM413 | BP | *Serratia plymuthica* | - | 3.4 ± 5.9 | 0.0 ± 0.0 |  |  |  |  |  |  |  |
| KMM414 | BP | *Micrococcus luteus* | - | 0.0 ± 0.0 | 0.0 ± 0.0 |  |  |  |  |  |  |  |
| KMM415 | BP | *Micrococcus antarcticus* | + | 0.0 ± 0.0 | 0.0 ± 0.0 |  |  |  |  |  |  |  |
| KMM416 | BP | *Micrococcus antarcticus* | + | 0.0 ± 0.0 | 0.0 ± 0.0 |  |  |  |  |  |  |  |
| KMM417 | BP | *Micrococcus luteus* | - | 0.0 ± 0.0 | 0.0 ± 0.0 | 0.013 ± 0.001 | 0.026 ± 0.005 | 0.046 ± 0.014 | 0.025 ± 0.006 | -0.036 ± 0.015 |  |  |
| KMM418 | BP | *Micrococcus luteus* | - | 0.0 ± 0.0 | 0.0 ± 0.0 | -0.004 ± 0.004 | 0.060 ± 0.032 | 0.012 ± 0.004 | -0.001 ± 0.002 | 0.019 ± 0.017 |  |  |
| KMM419 | BP | *Hafnia paralvei* | - | 55.8 ± 9.5 | 35.3 ± 0.0 |  |  |  |  |  |  |  |
| KMM420 | BP | *Micrococcus luteus* | - | 6.8 ± 5.9 | 0.0 ± 0.0 |  |  |  |  |  | 2.860 ± 0.125 | -0.007 ± 0.022 |
| KMM421 | BP | *Micrococcus luteus* | + | 14.7 ± 12.7 | 0.0 ± 0.0 | 0.023 ± 0.011 | 0.051 ± 0.007 | 0.012 ± 0.008 | 0.080 ± 0.042 | -0.006 ± 0.003 |  |  |
| KMM422 | BP | *Micrococcus sp.* | + | 34.0 ± 33.4 | 0.0 ± 0.0 |  |  |  |  |  |  |  |
| KMM423 | BP | *Micrococcus luteus* | + | 0.0 ± 0.0 | 0.0 ± 0.0 | 0.031 ± 0.005 | 0.009 ± 0.003 | 0.064 ± 0.071 | 0.014 ± 0.004 | -0.032 ± 0.023 |  |  |
| KMM424 | BP | *Micrococcus luteus* | + | 0.0 ± 0.0 | 0.0 ± 0.0 |  |  |  |  |  |  |  |
| KMM425 | BP | *Micrococcus luteus* | + | 0.0 ± 0.0 | 0.0 ± 0.0 |  |  |  |  |  |  |  |
| KMM426 | BP | *Micrococcus luteus* | + | 0.0 ± 0.0 | 0.0 ± 0.0 |  |  |  |  |  | 2.906 ± 0.177 | 0.009 ± 0.019 |
| KMM427 | BP | *Micrococcus sp.* | + | 0.0 ± 0.0 | 0.0 ± 0.0 |  |  |  |  |  |  |  |
| KMM428 | BP | *Micrococcus luteus* | + | 0.0 ± 0.0 | 0.0 ± 0.0 |  |  |  |  |  |  |  |
| KMM429 | BP | *Micrococcus sp.* | + | 0.0 ± 0.0 | 0.0 ± 0.0 |  |  |  |  |  |  |  |

| Supplementary Table 2. Top 21 isolates with the highest esterase activity on 1% tributyrin agar at 48 hours post-inoculation. | | | |
| --- | --- | --- | --- |
| Microbe ID | **Species** | **Population** | **Esterase Activity on 1% Tributyrin (mm^2^)** |
| KMM 191 | *Pseudomonas germanica* | Westchester Lagoon | 299.8 ± 81.4 |
| KMM 195 | *Pseudomonas trivialis* | Westchester Lagoon | 286.4 ± 31.4 |
| KMM 386 | *Aeromonas veronii* | Rabbit Slough | 276.7 ± 46.6 |
| KMM 387 | *Aeromonas veronii* | Rabbit Slough | 276.7 ± 46.6 |
| KMM 249 | *Pseudomonas sesami* | Mud Lake | 266.5 ± 47.0 |
| KMM 385 | *Aeromonas veronii* | Rabbit Slough | 236.9 ± 44.6 |
| KMM 230 | *Pseudomonas germanica* | Westchester Lagoon | 227.8 ± 47.6 |
| KMM 218 | *Aeromonas sobria* | Mud Lake | 226.2 ± 0.0 |
| KMM 367 | *Aeromonas veronii* | Rabbit Slough | 226.2 ± 0.0 |
| KMM 364 | *Aeromonas veronii* | Rabbit Slough | 217.0 ± 15.9 |
| KMM 376 | *Aeromonas veronii* | Rabbit Slough | 217.0 ± 15.9 |
| KMM 214 | *Aeromonas sobria* | Mud Lake | 215.2 ± 97.1 |
| KMM 366 | *Aeromonas veronii* | Rabbit Slough | 199.2 ± 26.7 |
| KMM 380 | *Aeromonas veronii* | Rabbit Slough | 199.2 ± 26.7 |
| KMM 379 | *Aeromonas veronii* | Rabbit Slough | 181.9 ± 29.0 |
| KMM 217 | *Aeromonas salmonicida* | Big Lake | 181.4 ± 14.9 |
| KMM 365 | *Aeromonas veronii* | Rabbit Slough | 181.4 ± 14.9 |
| KMM 377 | *Aeromonas veronii* | Rabbit Slough | 181.4 ± 14.9 |
| KMM 270 | *Pseudomonas azotoformans* | Cheney Lake | 173.3 ± 25.2 |
| KMM 293 | *Chromobacterium aquaticum* | Cheney Lake | 173.3 ± 25.2 |
| KMM 301 | *Pseudomonas protegens* | Cheney Lake | 173.3 ± 25.2 |
